# Rapid adaptation in large populations with very rare sex: scalings and spontaneous oscillations

**DOI:** 10.1101/233320

**Authors:** Michael T. Pearce, Daniel S. Fisher

## Abstract

Genetic exchange in microbes and other facultative sexuals can be rare enough that evolution is almost entirely asexual and populations almost clonal. But the benefits of genetic exchange depend crucially on the diversity of genotypes in a population. How very rare recombination together with the accumulation of new mutations shapes the diversity of large populations and gives rise to faster adaptation is still poorly understood. This paper analyzes a particularly simple model: organisms with two asexual chromosomes that can reassort during rare matings that occur at a rate *r*. The speed of adaptation for large population sizes, *N*, is found to depend on the ratio ~ log(*Nr*)/log(*N*). For larger populations, the r needed to yield the same speed deceases as a power of *N*. Remarkably, the population undergoes spontaneous oscillations alternating between phases when the fittest individuals are created by mutation and when they are created by reassortment, which—in contrast to conventional regimes—decreases the diversity. Between the two phases, the mean fitness jumps rapidly. The oscillatory dynamics and the strong fluctuations this induces have implications for the diversity and coalescent statistics. The results are potentially applicable to large microbial populations, especially viruses that have a small number of chromosomes. Some of the key features may be more broadly applicable for large populations with other types of rare genetic exchange.

## 1. Introduction

The reasons for the ubiquity of sex, or more broadly genetic exchange, across all domains of life has been the subject of a long and ongoing debate (reviewed recently by 1 and 2). Evolutionary explanations for sex must explain its advantages over its disadvantages in a spectrum of evolutionary scenarios and justify the benefits of its continued maintenance. Producing and maintaining diversity for selection to act on is a traditional explanation for sex. But when genetic exchange is a rare process—as it is now for many groups of organisms—even analyzing such advantages quantitatively can be challenging. Between instances of genetic exchange, asexual evolution produces genetic correlations in the population that limit the utility of sex. Close relatives gain little by mating with each other, so the evolution of a population dominated by clones will depend sensitively on the how its diversity is created and maintained by the combined effect of asexual accumulation of mutations and genetic exchange.

In large populations—as microbial populations usually are—many beneficial mutations can occur each generation. In asexual populations these mutations compete with each other and only one can ultimately takeover the population: such competition is known as clonal interference [3]. A classic hypothesis for the advantage of sex is the Fisher-Muller effect: sex reduces the competition between different beneficial mutations by enabling them to recombine onto the same genome, thus decreasing the “wastage” of beneficial mutations [4, 5, 6, 7, 8, 9]. But to understand the Fisher-Muller effects quantitatively in populations with low rates of recombination, a detailed understanding of asexual evolution in large populations is needed. A major complication beyond the simple clonal interference picture is that multiple beneficial mutations can arise on the same lineage before any of them fix. In this case, the success of a new mutation strongly depends on the genetic background on which it arises: only on an already very fit background does it have a substantial chance of fixing [10, 11, 12, 13]. Evidence of clonal interference and complex accumulation of beneficial mutations has been found in many laboratory experiments with bacteria, viruses, and *S. cerevisiae* (14; 15; 16; 17), as well as in natural viral populations [18, 19, 20].

To understand the interplay between clonal interference, acquisition of multiple beneficial mutations, and recombination, analysis of simple scenarios and models is needed. If the environment has recently changed, many beneficial mutations can become available and, at least for some time, the population can evolve under sustained directional selection with the supply of beneficial mutations only slowly depleting. (This appears to be the case in the long-term experiments of Lenski and collaborators, see 21.) The simplest model is to assume that only beneficial mutations occur, that their supply is not depleted, that they all have the same fitness effect, s, and that these effects are additive. There is now a large body of work analyzing asexual evolution in this model (22, 23, 11, 24, 25; reviewed in 26). For small populations the evolution is mutation limited with a beneficial mutation arising occasionally and sweeping to takeover the population before another occurs: the rate of fitness increase—the *speed*, υ, of the evolution—is then simply proportional to the population size, *N*, times the beneficial mutation rate, *U* (= *U_b_*). But in large populations, multiple mutations arise, compete—via clonal interference—and accumulate mutations in the same lineage before any fix. This results in a broad fitness distribution that forms a traveling-wave moving steadily towards higher fitness with a speed that grows only logarithmically with *N*. The statistics of the genetic diversity in such rapidly evolving asexual populations have also been analyzed: the phylogenies are characterized by multiple mergers and skewed branching, and the site frequency spectrum is strikingly different than neutral theory [27, 28].

Simple models of rapid adaptation have been extended to include the effects of sex. With enormous recombination rates, all mutations are independent and clonal interference is negligible. But in large populations, with any reasonable recombination rate mutations still compete. Nevertheless, the behavior simplifies for relatively high recombination rates, in particular for facultative sexuals with recombination frequent enough that mutations have a substantial chance of recombining onto a good genetic background before they are out-competed to extinction. For this to occur, the recombination rate, *r*, needs to be comparable or larger than the selective strength, *s*, up to logarithmic factors [29]. In this regime, the dynamics of new mutations are determined by the distribution of fitness backgrounds on which they arise and with which they can recombine. When many beneficial mutations are segregating at the same time, linkage is very transient and the correlations between mutational frequencies are weak: the behavior can then be analyzed by following the statistics of single mutations. To fix, some of the descendants of mutations that arise need to recombine onto a high fitness background, grow in number, recombine again, and so on until the mutant population becomes large enough to avoid extinction by linkage to “only” average genetic backgrounds. Neher et al. [29] find that the speed of evolution grows linearly or quadratically (depending on the model) with the recombination rate, *r*, because more and more mutations can segregate in parallel.

For obligate sexuals with linear chromosomes, the behavior also simplifies somewhat. The chromosomes act roughly as if broken up into effectively asexual segments whose evolution can be approximated by rapid asexual adaptation within each of the weakly correlated segments with the effective recombination rate between these being comparable to the selection coefficients. The speed of evolution is then proportional to the genomic recombination rate (map length), *R*, times log(*N*) [30, 31, 29, 32].

We are interested in the behavior with much lower recombination rates—not only *r* ≪ *s*, when the previous analyses already breakdown, but down to when *r* is of order an inverse power of the population size. This regime is increasingly important for very large and nearly asexual populations since the dynamics become increasingly sensitive to small recombination rates for larger populations. Recombination is infrequent enough that clonal growth, asexual accumulation of multiple mutations, and close relationships between recombining genomes are essential. As well shall see, the behavior is rather more complex. Rouzine and Coffin [33, 34, 35] have studied the purging of deleterious alleles starting from standing variation when mating is very rare but still results in a lot of recombination when it does occur. They found that the interplay between exponential growth of subpopulations from selection and recombination to produce higher fitness genomes results in a rate of fitness increase that depends only logarithmically on the frequency of sex. They also considered the effects of correlations due to common descent, which are an important effect for rare sex [36], and approximated the effects of these [35]. But these studies do not consider a crucial feature: how diversity in the fitness distribution is created and maintained by an influx of new mutations. It is the interplay between mutation, selection, and recombination that needs to be understood.

Simulations of simple models of asexual evolution can be carried out efficiently. For many aspects only fitnesses are relevant, and aspects of the diversity, such as the site frequency spectrum of individual mutations, can be inferred from methods based on fitness classes [27, 28]. But simulations of sexual populations with low rates of recombination need to keep track of the full *genomic* diversity. This can be computationally prohibitive for large population sizes, such as those found in microbial experiments, because they require keeping track of a huge number of genomes (roughly of order NU). Thus studying simpler models that include the key effects of both asexual mutation accumulation and occasional recombination is called for.

In this paper, we study a particular—and in some ways the simplest—compromise model: a facultative sexual population with two asexual chromosomes that can undergo reassortment but not recombination within the chromosomes. Under an assumption of additivity of the fitnesses of the two chromosomes, only the fitnesses of each chromosome are needed. This model was introduced and studied by Park and Krug [37], who focused on the limit when mating is frequent enough that the speed saturates at its value for obligate sexuals. Here we focus on the rare reassortment regime and analyze the crossover from asexual to obligate sexual. Because mating is rare, we can leverage much of the intuition and results from the asexual case. Reassortment can provide many of the benefits of recombination, so genome segmentation and reassortment could have been important in the origin of sex [38].

Although not the primary motivation, we note that models with reassortment of chromosomes but no recombination within them are natural for some RNA viruses such as influenza that have segmented genomes [39]. When two viruses co-infect a cell, segments from both can be repackaged into new viral particles, which results in reassortment. The rate of reassortment thus depends on the probability of co-infections and hence on the ratio of viruses to host cells so that reassortment is rare when viral densities are low.

## 2. Model and Parameters

We consider a population of N haploid individuals with genomes consisting of two chromosomes. They are facultatively sexual and two individuals *mate at a rate r*, resulting in the reassortment of their two chromosomes but no recombination. *Beneficial* mutations occur at a rate U (= *U_b_*) per chromosome. We are interested in continual evolution of large populations for which beneficial mutations are the driving force of the dynamics and deleterious mutations have minor effects [11]. Thus we consider an infinite-sites model in which all mutations are beneficial and each chromosome can accumulate any number of such mutations with no back mutations.

For simplicity, we take all mutations to have the same small (log-)fitness effect *s* ≪ 1 and assume additive fitnesses. The fitnesses of the two chromosomes, *X* and *Y*, are then simply *s* times the number of mutations on each, and the total fitness is *Z* = (*X* + *Y*). Despite the assumption of fixed effect size, the basic results of our analysis should hold more generally: asexual evolution with a distribution of mutation sizes has been shown to be dominated by a small range around a single effective *s* and an effective *U* if the distribution of mutation sizes falls off quickly enough [40, 41].

When mating, parental chromosomes are reassorted. Parents with fitnesses (*X*_1_, *Y*_1_) and (*X*_2_, *Y*_2_) will produce offspring with (*X*_1_, *Y*_2_) and (*X*_2_, *Y*_1_). Individuals are chosen randomly to undergo reassortment. Therefore the probability of reassortment producing the genome (*X, Y*) is proportional to the number of individuals with *X* and the number with *Y*. With discrete values of fitness, the subpopulations of the “fitness classes” we denote *n*(*X, Y*) and the total number with *X* by *n*(*X*) = *∑_Y_ n*(*X,Y*). Longterm increases in fitness rely on the nucleation of new high fitness classes by either mutation or reassortment. The parameters introduced in this section and main variables used throughout the text are summarized in table 1.

**Table 1:**
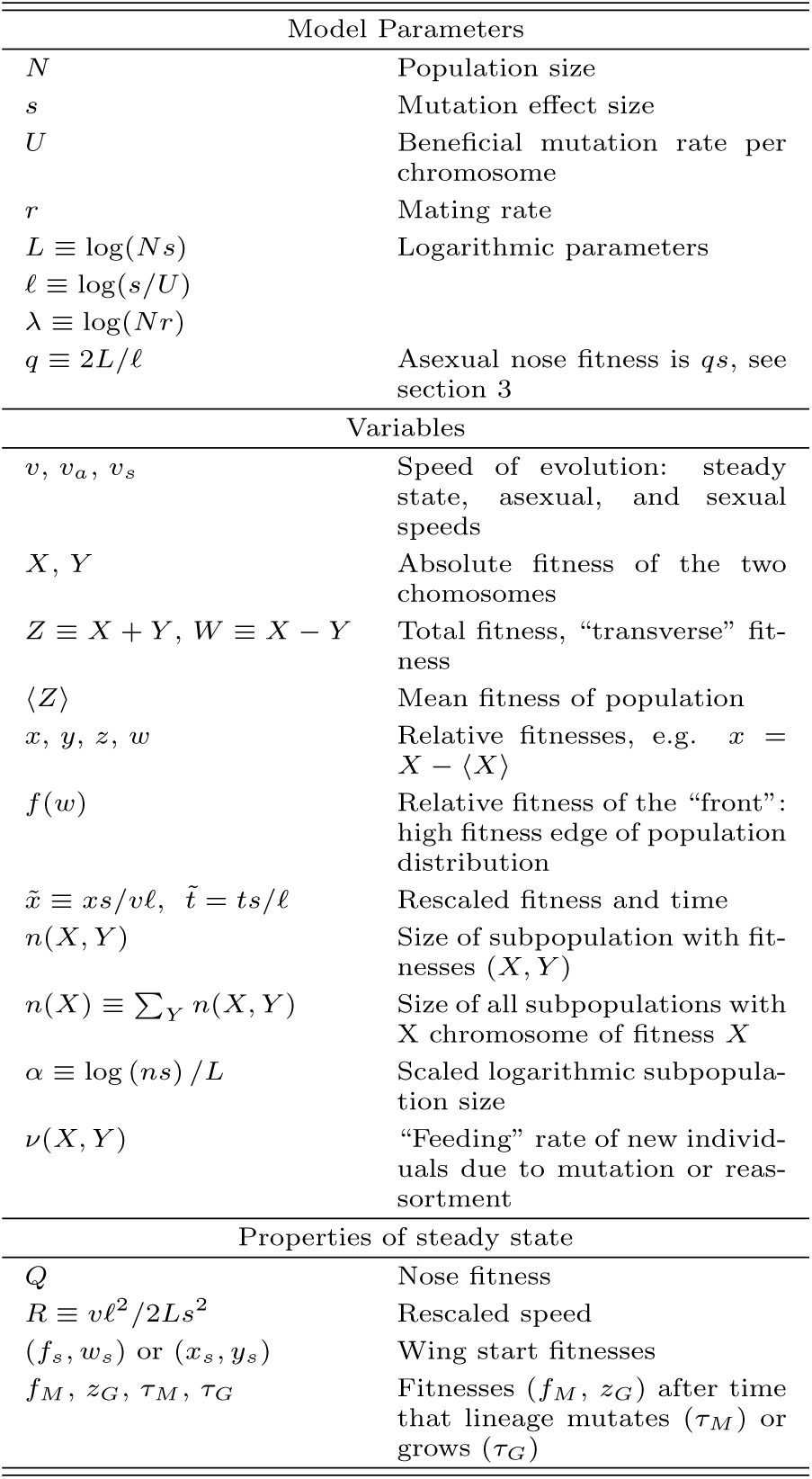
Parameters and variables.

| Model Parameters |  |
| --- | --- |
| $N$ | Population size |
| $s$ | Mutation effect size |
| $U$ | Beneficial mutation rate per chromosome |
| $r$ | Mating rate |
| $L \equiv \log(Ns)$ | Logarithmic parameters |
| $\ell \equiv \log(s/U)$ | |
| $\lambda \equiv \log(Nr)$ | |
| $q \equiv 2L/\ell$ | Asexual nose fitness is $qs$, see section 3 |
| Variables |  |
| $v, v_a, v_s$ | Speed of evolution: steady state, asexual, and sexual speeds |
| $X, Y$ | Absolute fitness of the two chromosomes |
| $Z \equiv X + Y, W \equiv X - Y$ | Total fitness, “transverse” fitness |
| $\langle Z \rangle$ | Mean fitness of population |
| $x, y, z, w$ | Relative fitnesses, e.g. $x = X - \langle X \rangle$ |
| $f(w)$ | Relative fitness of the “front”: high fitness edge of population distribution |
| $\tilde{x} \equiv xs/v\ell, \tilde{t} \equiv ts/\ell$ | Rescaled fitness and time |
| $n(X, Y)$ | Size of subpopulation with fitnesses $(X, Y)$ |
| $n(X) \equiv \sum_Y n(X, Y)$ | Size of all subpopulations with $X$ chromosome of fitness $X$ |
| $\alpha \equiv \log(ns)/L$ | Scaled logarithmic subpopulation size |
| $\nu(X, Y)$ | “Feeding” rate of new individuals due to mutation or reassortment |
| Properties of steady state |  |
| $Q$ | Nose fitness |
| $R \equiv v\ell^2/2Ls^2$ | Rescaled speed |
| $(f_s, w_s)$ or $(x_s, y_s)$ | Wing start fitnesses |
| $f_M, z_G, \tau_M, \tau_G$ | Fitnesses $(f_M, z_G)$ after time that lineage mutates ($\tau_M$) or grows ($\tau_G$) |

The schematic stochastic differential equation for the dynamics of a subpopulation (approximated as being continuous) is

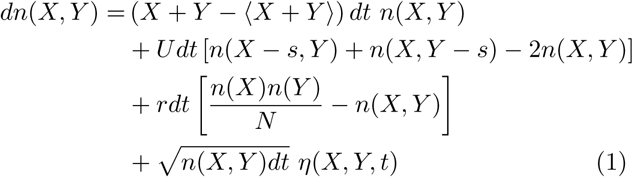

The first term represents growth and selection determined by the relative fitness above the population-mean fitness, 〈Z〉: henceforth we will denote absolute fitness (in units of s) with capital letters, e.g. *X*, and relative fitness with lowercase, e.g. *x* = *X* − 〈X〉. The second term is the net influx of mutations into the (*X, Y*) fitness class. The third term describes reassortment: the number of new offspring is proportional to the total number of matings in the population, *Nr*, times the frequencies of the two chromosomes with the needed fitnesses, i.e.. *n*(*X*)/*N* and *n*(*Y*)/*N*. The final term represents the stochasticity of births and deaths. The distribution of the random variables {*η*} are essentially independent gaussians for each fitness class with small—but essential—corrections to enforce the fixed population size constraint [40]. The 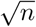 factor mimics the effects of discrete individuals (the “diffusion approximation”), and allows subpopulations to go extinct. We will approximate this stochasticity in other ways that do not affect any substantial properties.

Primary quantities of interest are the mean speed of evolution defined as

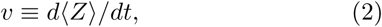

the shape of the two-dimensional distribution of non-zero sub-populations—in particular its fitter-than-mean boundary we call the *front*—and the fluctuations of these quantities.

We will consider the range of parameters for which the asexual dynamics simplify: strong selection (*N_s_* ≫ 1) and weak mutation (*s* ≫ *U*). This regime is applicable for large microbial populations. We are interested in the effects of multiple mutations that are simultaneously segregating in the population, and thus focus on the regime *NU* ≫ 1 in which many beneficial mutations occur each generation. (See, for example, [42] for the effects of sex when clonal interference is weak and NU is order one.) The roles of these assumptions will be discussed in section 3. Park and Krug [37] studied the limit *r* ≫ *s* of the two chromosome model. They found that in this regime the speed of evolution rapidly approaches the obligate sexual (*r* → ∞) limit. When *r* is substantially greater than s, the reassortment rate is high enough that any linkage between chromosomes is broken up before growth becomes important, so each chromosome evolves independently. Thus for *r* ≪ *s*, *υ*(*U*) ≈ 2*υ_a_*(*U*) where *υ_a_(U*) is the asexual speed for a single chromosome. Our focus is the regime when mating is rare (*r* ≪ *s*). In this regime almost all subpopulations grow clonally with negligible loss or gain due to reassortment. But reassortment is crucial when it produces individuals in new fitness classes earlier than mutations can. As the mating rate is increased from zero to much larger than *s*, *v* should increase from *υ_a_*(2*U*) to 2*υ_a_*(*U*). It is this crossover regime, in which the interplay between mutation accumulation and reassortment is most subtle, that we particularly wish to understand.

Because the subpopulations with fitnesses higher than the mean are growing exponentially, the times at which key events occur depend only logarithmically on the parameters. Thus it is useful to define logarithmic variables for the large parameters of the model:

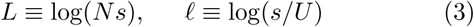

As in the asexual case reviewed below, the behavior and analysis of the model simplifies in an asymptotic regime where the logarithmic parameters are themselves large, and more so if in addition *L* ≫ *ℓ* ≫ 1. Although these are never strong inequalities in practice, the approximations they lead to capture the qualitative behavior and are quite good quantitatively for reasonable parameter ranges [11].

With reassortment, new high fitness subpopulations are produced by subpopulations that are growing exponentially. Thus again characteristic times might be expected to depend only logarithmically on parameters. It is convenient to define a logarithmic measure for the reassortment rate in terms of the population-total rate of mating:

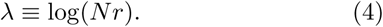

As the reassortment rate increases from 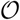(1/*N*) to 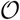(*s*), λ increases from 0 to *L*. (When λ < 0 there are typically no reassortments in the whole population and their effect is negligible.)

For very large populations an important simplification is that many sums over subpopulations are dominated by a small subset of them. Subpopulations near the boundary of the fitness distribution have sizes, *n*, of order unity while those near its maximum have sizes a substantial fraction of *N*. Thus log(*ns*) ranges from 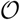(1) to *L*. The exponential growth (or decay) with time implies that it is natural to express these using a normalized logarithmic variable α such that *ns* ≡ exp(*Lα*). A sum over subpopulations can then be approximated as∫ exp(*Lα*) ≈ exp(*Lα*_max_) for *L* ≫ 1, where we have dropped factors that are not exponential in *L*. For example, the sum *n*(*X*) = *∑_Y_ n*(*X, Y*) determines how many *X* chromosomes are available for reassortment. For large *L*, nearly all such *X* chromosomes come from one subpopulation or from a narrow range of subpopulations relative to the full width of the fitness distribution.

## 3. Review of Asexual Dynamics

In the limit of no mating, the dynamics are asexual. Qualitative concepts and scalings from the asexual dynamics are crucial for understanding the regimes of rare mating, thus we first review the asexual limit following the heuristic analysis of Desai and Fisher [11]. The picture of asexual evolution can be greatly simplified by a separation of scales. The growth of a clonal subpopulation is affected by both stochastic fluctuations and the nonlinear population size constraint. However, for strong selection (*Ns* ≫ 1) stochastic fluctuations only matter when a subpopulation is relatively small, long before it grows enough that the nonlinear constraint becomes important. Conversely, for the large subpopulations that dominate, the total population size constraint is crucial but the fluctuations are small. This separation of scales allows the influence of fluctuations and the population size constraint to be analyzed separately. But an essential property of the dynamics is the coupling between the fluctuations of small subpopulations and the effects of these fluctuations when those subpopulations become large and nonlinearities become important: this is what makes the full stochastic dynamics so difficult to analyze theoretically [40].

The crucial subpopulation is the one with highest fitness, which on average grows exponentially faster than the others. Conditional on a new subpopulation surviving drift and becoming established, its fluctuations can be neatly packaged into an establishment time, roughly the time at which the lineage appears to have started growing deterministically. A lineage with fitness *z* above the mean has probability *z* of establishing. At long times, an established lineage grows as

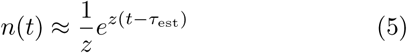

where the random variable τ_est_ is the establishment time whose distribution captures the randomness of the early-time fluctuations of a new sub-population [11].

In the multiple mutations regime (*NU* ≫ 1), there are many subpopulations that are fitter than the mean. Mutations from the fittest subpopulation—referred to as the *nose* of the distribution—will start nucleating a new fitter subpopulation. This new nose has some fitness, *Q*, above the mean. This subpopulation, *n_Q_*, receives mutations from the subpopulation *n_Q−s_*. In the weak mutation limit (*s/U* ≫ 1), *n_Q−s_* will already be growing deterministically before the nose is likely to establish. After establishing with size 1/*Q*, the subpopulation grows as *n_Q−s_* = exp [(*Q* − *s*)*t*] /*Q* with *t* measured from its own establishment time. It can be shown that the establishment time of *n_Q_* has a mean of

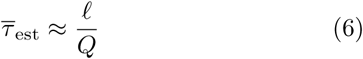

for large *ℓ* [11].

For rapid asexual adaptation, the balance of selection and mutation results in a steady state population distribution whose mean fitness increases at some average speed *υ_a_*. For consistency, the nose must advance, one mutation at a time, at the same speed *υ_a_*. Hence *υ_a_* = *s*/τ̅_est_.

Behind the nose, the dynamics are essentially deterministic with negligible contributions from new mutations (for *ℓ* ≫ 1). The speed and shape of the distribution are determined by the balance between selection and the constraint on the total population size. After establishment, the subpopulation formerly at the nose will grow deterministically but with ever decreasing relative fitness, *Q* − *υ_a_*. This subpopulation grows until its fitness is the same as the mean and reaches a maximum size in a time τ_nm_ = *Q*/*υ_a_*, called the nose-to-mean time. At this time, it has become the largest subpopulation with a size of order *N*. Thus (to logarithmic accuracy) this subpopulation size is

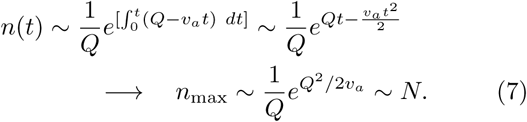

Thus for consistency we need *υ_a_* = *Q*^2^/2log(*Ns*) where we have dropped a factor of *Q*/*s* inside the logarithm since log(*Ns*) ≫ log(*Q/s*). Equating this speed to the advance of the nose, *υ_a_* = *s*/τ̅_est_ ≈ *Q_s_*/*ℓ*, yields

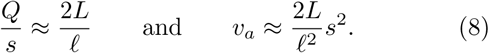

Note that the speed does not depend directly on the total mutation rate *NU* because of clonal interference between the mutations that arise. Only the small fraction of mutations that arise near the anomalously fit backgrounds near the nose have a chance of fixing. The shape of the fitness distribution also follows simply by tracking the subpopulations: it is very close to gaussian except right at the nose where it is “cutoff” [22] and strongly fluctuating [40].

### 3.1. Approximations and their validity

The accuracy of the approximations that lead to eq. (8) depend on combinations of parameters. The basic regime of validity of the primary approximations is *L* → ∞, but how large *L* must be depends also on *ℓ* and differs somewhat depending on the quantity of interest. If the number of mutations of the nose above the mean, 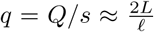, is only a few, then there are several sources of corrections to the simple asymptotic expressions. The nose fitness changes from *Q* to *Q* – *s* while establishing, so a somewhat more accurate analysis using the average fitness 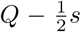 yields *υ_a_*/*s*^2^ ≈ (2*L* − *ℓ*)/*ℓ*^2^. This also matches correctly to the speed at the crossover, at *L* = *ℓ*, from the successive mutations regime with *υ_a_* ≈ *NUs*^2^ for *NU* ≪ 1/*ℓ* to the multiple mutations regime for larger NU. However this assumes smooth motion of the mean, which is not the case for small *q*. The motion is jerky even in the deterministic approximation and the fitness distribution is not so well approximated by a smooth gaussian. Smooth motion of the mean requires high enough speeds that the root mean square width, 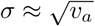, of the fitness distribution is greater than a single mutation, *s*: this requires *L* ≳ *ℓ*^2^, or *q* ≳ *ℓ*. Note, however, that because sums over the discrete populations approximate very well a smooth gaussian, in practice already for *q* ~ *ℓ*/3 the effects of the jerkiness are small [40]. Moreover, for many properties of interest, the effects of the jerky motion of the mean are minor as long as q is relatively large.

We will generally analyze the asymptotically large *L* limit in which most aspects of the asexual dynamics become essentially deterministic and the fitness distributions smooth enough that the fitness can be considered as a continuous variable. There are major simplifications that we make use of when *ℓ* is also large. We largely restrict the analysis to this limit and discuss corrections when needed to compare with simulations and study crossover regimes. A discussion of the effects of fluctuations and their scaling with the reassortment rate can be found in section 8. As we shall see, much of the behavior is captured, even quantitatively, for realistic values of parameters that are far from the asymptotic limits of validity of the analyses.

### 3.2. Scaled variables

The asexual analysis suggests some basic rescalings that will be useful for studying the two chromosome model. The important fitness and time scales are the nose fitness *Q* = *υ_a_ℓ*/*s* and the nose-to-mean time, τ_nm_ = *Q*/*υ_a_* = *ℓ*/*s*. Thus a natural set of rescalings for fitness and time, which we will use more generally, are

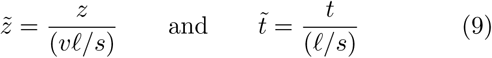

where *υ* is the *actual* average speed—as yet unknown with reassortment but in general greater than the asexual speed *υ_a_*. In these rescaled variables the speed is unity and a subpopulation’s fitness relative to the advancing mean has a simple expression:

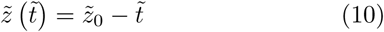

For the two chromosome model, we will define

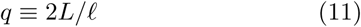

for convenience. In the asexual limit *q* is the number of mutations that the nose is above the mean, eq. (8). While this is *not* true for nonzero mating rates, *q* is a convenient parameter combination that turns out to be always within a factor of two of the number of mutations between the mean and the nose. The asymptotic regime in which most of our analyses become accurate corresponds to *q* ≫ 1, so that many fitness classes are populated and the fitness is effectively continuous.

## 4. Simulation Results

The separation of scales when evolution is rapid allows efficient simulations for large population sizes. In particular, one can ignore stochasticity except in small subpopulations for which fluctuations are important. Although these fluctuations will eventually effect the large subpopulations, we directly incorporate the resulting nonlinearities by keeping the total population size fixed. The establishment size (above which it is unlikely to go extinct) for a subpopulation with fitness *z* is *n*_est_ ~ 1/*z*, so we conservatively set a threshold for stochasticity of *n* < 10/s. Eq. 1 is used to generate the expected number, *n*_exp_, of individuals for each subpopulation (*X, Y*) for the next time step. For *n*_exp_ ≥ 10/*s* we simply set *n* = *n*_exp_, corresponding to deterministic growth. For small subpopulations *n*_exp_ < 10/*s*, we sample n from a Poisson distribution with mean *n*_exp_. This scheme slightly alters the birth-death process but the differences are small even on linear scales and are negligible when the logarithmic parameters that control the very large *N* behavior become substantial. (In an exponentially growing population the fluctuations are dominated by early times: from a size *n*_0_, the late time fluctuations in the subpopulation size are 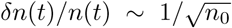. With *n*_0_ = 10/*s* these fluctuations are at most a few percent, much smaller than those from the time when the subpopulation is establishing, when *n*(*t*) ~ 1/*qs* or smaller, that are accounted for by our simulation scheme.)

To understand the behavior, it is useful to study a fully deterministic approximation to the dynamics, For the asexual case, a deterministic approximation of the nose dynamics approximates very well the mean shape and speed of the fitness distribution. This approach of using a deterministic model with a cutoff at high fitness goes back to the study of the asexual model by Tsimring et al. [22]. What is needed is a way to decide when a new deterministic subpopulation establishes. One simple approach is based on the expected number of established lineages. Consider an influx *ν*(*t*) of new mutants into an initially empty sub-population with fitness *z*. Then the expected number of established lineages at time *t* is

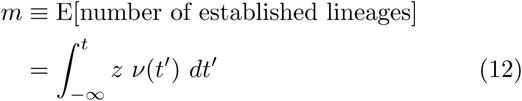

For our specific deterministic approximation, whenever *m*(*t*) reaches an integer, we add a new lineage of size 1/*z* to the subpopulation. This procedure removes stochasticity without allowing fractions of an individual to have an effect. For the asexual case, it corresponds almost exactly to the approximation used above in section 3 of treating the stepward advance of the nose at constant intervals of τ̅_est_.

An implementation of the stochastic and deterministic simulations used in this paper are provided via an online data repository, see [43]. Figure 1 shows results for the mean speed from the stochastic simulations and compares these to steady state approximations analyzed in the following sections. The quantity plotted is the ratio of the increase in speed above the two chromosome asexual speed, *υ* − *υ_a_*(2*U*), to the increase of a fully sexual population, *υ_s_* − *υ_a_*(2*U*) (with reassortment but no recombination). This ratio thus goes from 0 to 1 as *r* is increased from 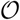(1/*N*) to 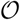(*s*). A primary result from the *steady state approximation* analyzed below is that, asymptotically, the speed should depend on the recombination rate only through the combination, λ/*L* ≡ log(*Nr*)/log(*Ns*). The simulation results show that the λ/*L* dependence indeed captures the overall scaling of the speed. In the asymptotic limit of large *ℓ* and large *q* = 2*L*/ *ℓ*, the normalized speed predicted for the steady state does not depend on *q* or *ℓ*. The stochastic simulations in fig. D.13 show that, indeed, the speed does not depend much on *q*, but at low reassortment rates there is substantial dependence on *ℓ*. We will elucidate and expand on various aspects of fig. 1 in the following sections, including the derivation of the steady state approximations and a discussion of the more complicated—and interesting—features of the dynamics beyond the mean speed.

**Figure 1:**
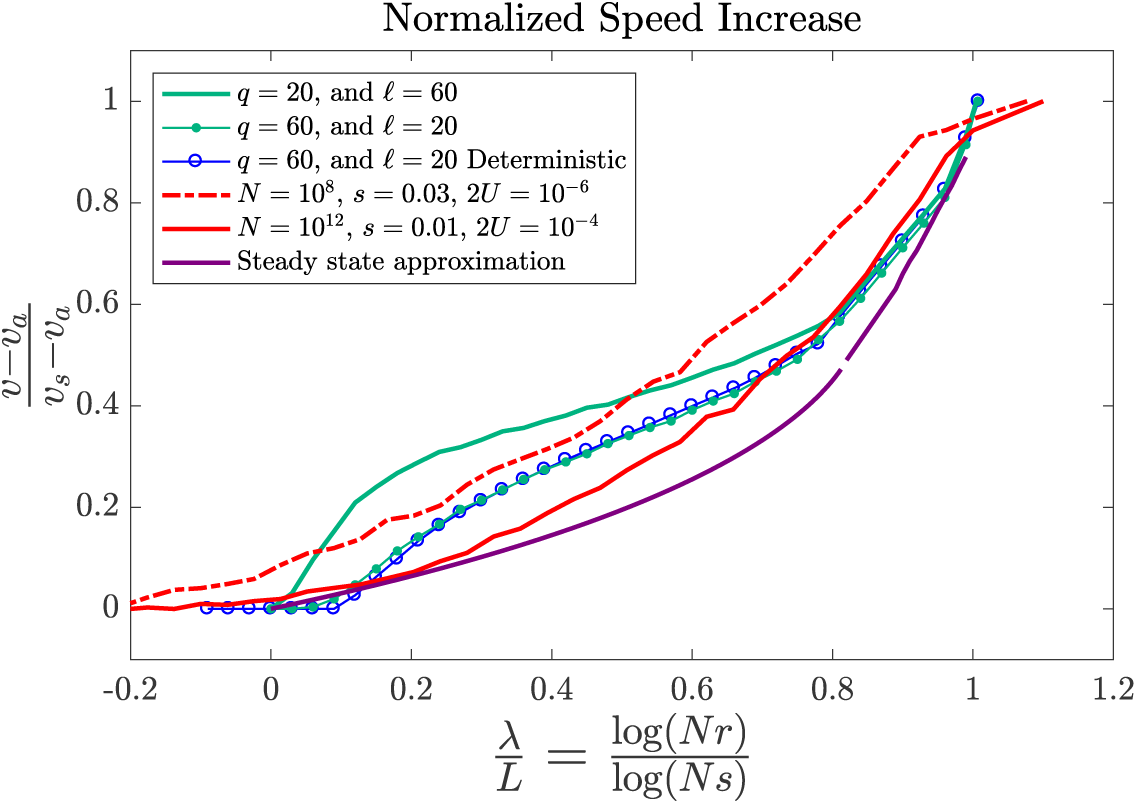
Stochastic simulation results for the speed of evolution for different values of *N*, *r*, and *U* compared to the steady state analysis and deterministic simulations. Simulations were run for a time 200 *ℓ*/*s* for the *q* = 20 and *q* =60 curves (green) and a time 600 *ℓ*/*s* for the *N* = 10^8^ and *N* = 10^12^ curves (red), with *ℓ*/*s* being roughly the time for the nose population to become the largest subpopulation. The quantity plotted is the normalized increase in speed due to reassortment: the ratio of the speed increase from asexual, *υ* − *υ_a_*, to the maximum speed increase in the sexual limit, *υ_s_* − *υ_a_*. The *q* = 20 and *q* = 60 curves are for asymptotically large populations, both with *L* ≡ log(*Ns*) = 600, so that the fitness distribution is essentially continuous. The red curves illustrate two realistic sets of parameter values for large microbial populations feasible in evolution experiments (These correspond to *q* ≅ 2.7, *ℓ* ≅ 11 for *N* = 10^8^ and *q* ≅ 9, *ℓ* ≅ 5.3 for *N* = 10^12^) The *N* = 10^8^ curve is shifted due to corrections of order one in the effective λ caused by fluctuations as discussed in section 5.1. The purple curves show the results of the steady state analysis which exhibits two regimes: low mating (λ/*L* < 0.8) and intermediate mating (λ/*L* > 0.8). The observed speed from the stochastic simulations also changes non-smoothly near the boundary between these two regimes. The blue curve shows the results of deterministic simulations illustrating excellent agreement with the stochastic simulations for a particular set of parameter values in the large *q* regime. A more extensive comparison of deterministic and stochastic simulations is in fig. D.13.

## 5. Two Chromosome Asexual Limit

Without reassortment, the two chromosome model is equivalent to asexual evolution but with each mutation having a label: X or Y. With twice the overall mutation rate, the speed is *υ_a_*(2*U*) ≈ 2*L*/log(*s*/2*U*)^2^ = 2*L*/(*ℓ* − log(2))^2^. (Note that the small log(2) correction to *ℓ* yields corrections to υ of the same order as other 1/ *ℓ* terms neglected in the approximations used in deriving the asexual speed, but in ratios its effects are noticeable for any reasonable *ℓ*.) In principle, the population distribution *n*(*X, Y*) could be inferred from the asexual diversity statistics derived in Desai et al. [28] by accounting for the random X or Y labels. We shall see that the subpopulations important for small reassortment have a very asymmetric division of mutations—anomalously high fitness on one chromosome and low or average fitness on the other— corresponding to tails of the distribution of labels. Thus to understand the behavior with reassortment we must track the dynamics of anomalous lineages that mutate mostly on a single chromosome. These are not so readily amenable to the asexual approaches of Desai et al. [28], thus we instead use a fitness class approach similar to section 3.

With two chromosomes there is a one-dimensional *front* in fitness space that marks the leading edge of the fitness distribution. A useful variable to parametrize the fitness distribution is the difference in fitness between the two chromosomes, *w* ≡ *X* − *Y* − 〈*X* − *Y*〉 = *x* − *y*, which on average does not change as the mean fitness advances. The front is the set of subpopulations with the highest fitness for a given *w* value and their fitnesses relative to the mean we denote *f*(*w*). These subpopulations at the front are the ones that have most recently established. The steady state asexual fronts are plotted in fig. 2 for different mutation rates. If the transverse coordinate, *w*, is integrated over, the nose of the resulting fitness distribution is dominated by only part of the front, and likewise the mean by only a small part of the range of *w*. Nevertheless, we shall see that with reassortment, a wider range of both the front and the bulk of the distribution will be important.

**Figure 2:**
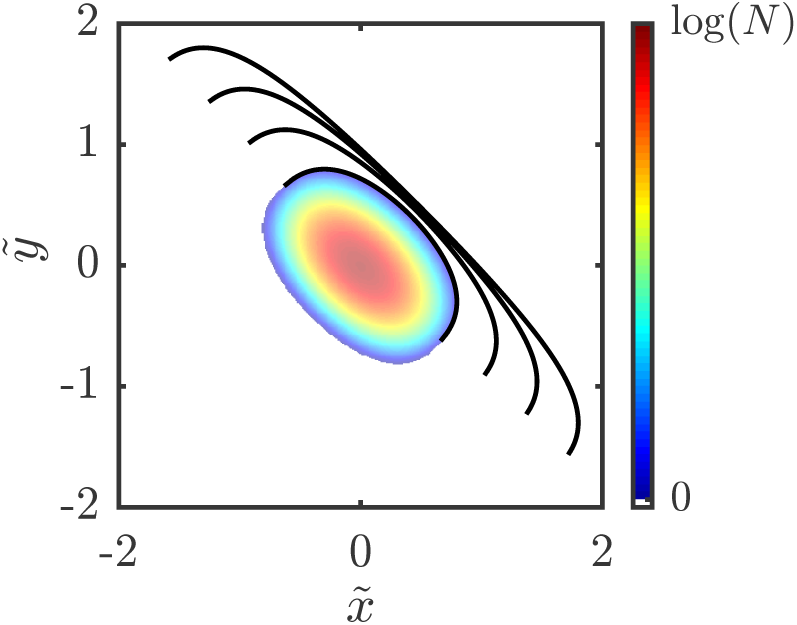
Shape of asexual fitness front *f* (*w*)—the boundary of the populated region of fitnesses, (*x,y*) of the two chromosomes—for several values of *ℓ* ≡ log(*s/U*): from inside to outside, *ℓ* = 4, 8, 16, and 32. The diagonal directions are the total fitness, *f* = *x* + *y* and the difference between the fitnesses of the two chromosomes, *w* ≡ *x* − *y*. The width of the distribution (in *w*) grows as ~ log(*ℓ*) for large *ℓ*. The colored region is the population distribution for *ℓ* = 4 in the deterministic approximation with colors indicating contours of log-population size. The population evolves by moving at constant speed in the upper right direction. Fitnesses are measured relative to the mean in normalized units.

The establishment time calculations in section 3 can be used to derive a self-consistency condition for a steady state front. In the limit of large *q* and *f* ≫ *s*, one can approximate the front as continuous, so that the condition will become a differential equation for *f*(*w*). A new mutant with (*X, Y*) can come from two parental fitness classes: (*X* − *s, Y*) and (*X, Y* − *s*). The combined size of the two feeding subpopulations is

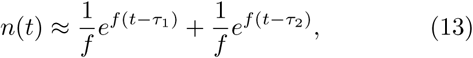

where τ_1_ and τ_2_ are their two establishment times (here ignoring the difference between *f* and *f* − *s* as in the asexual case). The combined subpopulation grows exponentially as exp[*f* (*t* − τ_0_)]/*f* with an effective establishment time

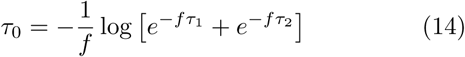

Therefore, we can directly use the result of eq. (6) to find the mean establishment time of the (X, Y) population:

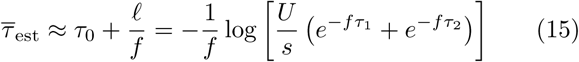

If one population never established (τ_2_ → ∞) we recover the one chromosome result, τ_est_ = τ_1_ + *ℓ*/*f*.

If we know the establishment times of all subpopulations along the front, then we can use eq. (15) to determine how the front advances with time. In the pseudo-deterministic approximation, as used in section 3, we expect a steady state shape for the front with the mean and front advancing at the same speed *υ* = *υ_a_*(2*U*). There is, of course, a trivial solution with a completely straight front, but this is inconsistent with the fixed population size constraint. Thus the steady state fitness distribution, and hence the front, must be localized in some range of *w*. The natural Ansatz is that the front has a single point (or two neighboring points) with maximum fitness: we will again call this point the *nose*. In two steps, each taking time *s/υ*, the nose will advance from (*X, Y*) to (*X* + *s*, *Y* + *s*).

The detailed analysis of the steady state front is in Appendix A: we summarize here its key features. When *q*, which characterizes the range of occupied fitness classes, is very large, we expect that the width of the distribution will be also. In this limit, the front can be approximated as a smooth curve. The front is comprised of two regions characterized by how establishment occurs. In the middle region around the nose, establishment occurs because of mutations from both parent populations. This means that τ_1_ and τ_2_ are close. In the outer “wings”, establishment is dominated by mutations from only one parent population because the other parental subpopulation established too late to contribute many mutants. This occurs when the difference in establishment times, |τ_1_ − τ_2_|, is much larger than the time, 1/*f*, for the populations to grow significantly.

The behavior in the wings is easier to understand and will be needed for the analysis with reassortment. Subpopulations at the front that are further from the nose have smaller fitness. So for the X wing, an X mutation from the X parent will establish earlier and with greater fitness than a Y mutation from the Y parent, thus establishment is mainly due to mutations from the X parent. Lineages in the X wing therefore stay at the front by accumulating mutations predominantly on the X chromosome, moving their descendant lineage even further from the nose. Such a lineage accumulates mutations more slowly than the nose advances so it loses relative fitness over time. Its total fitness is *f* = *X* + *Y* − *υt* with *Y* constant. Since a mutation of size *s* is added to the X chromosome in a time *ℓ*/*f*, *X* increases at speed 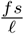. Therefore

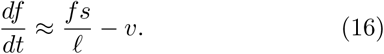

At the same time, *w* = *X* − *Y* is increasing at rate 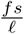. For the shape of the front to remain the same requires 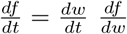, which yields a differential equation for *f*(w) valid in the X-wing. The shape in this region does not depend on any parameters, since eq. (16) becomes 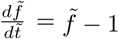 in our normalized variables.

In the middle region near the nose, the behavior is more subtle: here the difference between *ℓ* and *ℓ* − log 2 (i.e. *U* and 2*U*) is important (the nose fitness satisfies a similar differential equation to 16 with *ℓ* replaced by *ℓ* − log2). The shape, *f*(*w*), of the steadily moving front near the nose thus satisfies a more complicated differential equation. Matching together this nose region with the X and Y wings, as described in Appendix A, yields a shape of the fitness distribution controlled entirely by *ℓ* with the other parameter *q* ≡ 2*L*/*l* only setting its overall size in fitness space. The shape of the front for different *ℓ* values is shown in fig. 2. The ratio of the width *w*_max_ to the max fitness *f*_max_, or “aspect ratio”, depends weakly on *ℓ*, going as

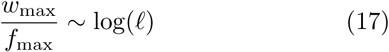

for large *ℓ*. Since *ℓ* ≡ log(*s/U*) is itself a logarithmic parameter, the dependence on the underlying parameters is very weak—although it will turn out to account for the *ℓ* dependence of the speed seen in fig. 1 at low λ/*L*.

Similarly to the aspect ratio, the time to relax to the asexual steady state depends weakly on *ℓ*. For our later discussion of oscillations, the relevant initial condition is a distribution with a nose fitness roughly *Q* but with the shape and width of the distribution far from the steady state, being substantially more curved near the nose. Tracking how the wings mutate away from the nose and lose fitness, one finds that the distribution relaxes to the steady state in a time 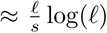 which can be several times the nose-to-mean time, *ℓ*/*s*.

Note that the analysis above and in Appendix A is formally valid in the limit in which *q* → ∞ *before ℓ* → ∞. This limit ensures that the thickness in f of the nose region is large compared to s (which requires *q* ≫ *ℓ*) so that the continuum approximation is justified. But the simulations show that the behavior is essentially the same in the opposite limit. In that regime, the mean as well as the front moves jerkily. The fitness does not vary smoothly across the front, but the timings of the establishments are still a smooth function of *w* so the continuum approximation can still be valid.

### 5.1. Initial deviations from asexual

In the very rare sex regime, as r is increased from zero the reassortment alters the asexual dynamics when it can result in the establishment of a new subpopulation in a fitness class beyond the front. The reassortment feeding has the form 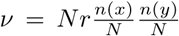 and establishment occurs when *ν* ~ 1, as detailed in section 6.1. For the asexual steady state analyzed above, there is a gently curving ridge of large subpopulations with fitness *z* = 0. In the approximation that those that can first reassort to the nose have *n*(*x*) ~ *n*(*y*) ~ *N* — the maximum possible — the condition for reassortment to increase the speed is simply λ/*L* = 0. As derived in appendix Appendix C, the curvature of the ridge, which arises from the curvature of the asexual front, reduces the size of the mating subpopulations and changes the borderline condition to λ/*L* ≈ 2/ *ℓ* for large *ℓ* consistent with the deterministic simulations in fig. D.13. However the stochastic simulations in fig. 1 show that the speed starts to increase already for λ/*L* ≅ 0. This is a consequence of fluctuations. Anomalously early establishments in the nose region of the front can result later in a temporarily broader population distribution that has large subpopulations sufficiently far apart to advance the nose via reassortment for smaller λ/*L*. In the asymptotic limit, these establishments due to fluctuations still require λ/*L* ≥ 0. But for modest size populations, deviations from asexual behavior before λ < *L* can occur, as seen in fig. 1 for *N* = 10^8^. For that population size, with *L* ≈ 15, the sub-exponential prefactors in *n*(*x*) and the feeding rate, *ν*, required for establishing a subpopulation are not negligible. The leading corrections can be incorporated into an effective reassortment parameter, λ_eff_ = λ + 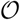(1). This will shift the speed curve by 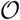(1/*L*), consistent with the simulations in fig. 1.

## 6. Reassortment Steady State

We now turn to an analysis of the behavior for small reassortment under the *assumption* of a steady state fitness distribution. This is a good starting point for understanding the key aspects of the dynamics and dependence on the basic parameters as well as analyzing fluctuations (as for the asexual model, see 40). As we shall see, for the two chromosome model, the steady state turns out to be unstable to oscillations as discussed in a later section. But it is still instructive to study the steady state because the important features of the oscillations—a cycle of accumulating mutations, clonal growth, and then reassortment— already occur in the steady state.

We make a basic *Ansatz* for the nature of the fitness front: that it has two types of regions that depend on how new subpopulations are nucleated at the front. In one region, the *reassortment front*, individuals in a new subpopulation are produced by reassortment of parents in the interior of the fitness distribution. In the other regions, called the *mutation wings*, individuals are the product of mutation and have a single parent that is (or more accurately, recently was) at the front. The wings are similar to those of the asexual limit already analyzed above.

The reassortment front is located in the middle of the front where *x* ≈ *y*, with the nose at *x* = *y* = *Q*/2 with the largest relative fitness, *Q*. For this region, parents typically have one quite high fitness chromosome and one quite average fitness chromosome, resulting in offspring that have two quite high fitness chromosomes. There is a mutation wing on either side of the reassortment front. Establishment in the wings is due to mutation because reassortment produces too few individuals. For large populations we expect the transitions between the reassortment front and the wings to be sharply delineated, i.e. the influx of individuals will either be dominated by reassortment or mutation.

The total population is dominated by the subpopulations that arise near the nose that later found near the peak of the distribution at (0,0). These subpopulations reach a maximum size after growing from a time *t* = *Q*/*υ*. Ignoring sub-exponential prefactors, the maximum size should be ~ *N* so

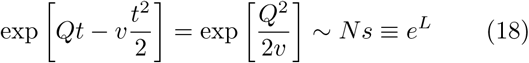

as for the asexual case. In terms of the scaled variable,

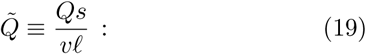

this implies that the the scaled speed is related to the scaled nose fitness by

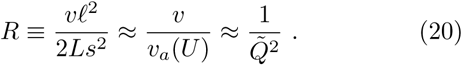

Using its relationship to 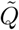, the speed drops out of the analysis of the fitness distribution’s shape, which then amounts to finding 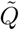 as a function of λ/L. There is a dual meaning of 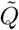: since the speed at which the nose can advance by mutations is *Qs*/ *ℓ*, 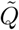 is the ratio between this would-be mutational-driven speed and the actual speed with reassortment. Alternatively, 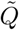 can be viewed as the nose-to-mean time in rescaled variables.

The mutation wing and the asexual front in section 5 share the same mutational dynamics and therefore obey the same differential equation for their shape, which depends on *ℓ*. In the large *ℓ* limit, however, the dynamics simplify because the wing will start with low enough fitness that only mutation from a single parent population matters. The shape of the mutation wing would then be independent of *ℓ*. We will discuss when this approximation is valid.

### 6.1. Low Mating Rate Steady State

At low mating rates, the reassortment front is narrow and is solely the product of mating between subpopulations that descend from the two mutation wings. These subpopulations typically have one anomalously fit chromosome and one close-to-average chromosome. The subpopulations along the reassortment front will grow to large sizes, but they are not important for further reassortment. Their fitness is split equally between the two chromosomes so they do not contribute especially fit chromosomes during mating.

The low mating steady state is relatively simple because the mutation wings are determined entirely by the boundary point between the two regions, called the *wing start*. We will solve for the steady state moving at a speed v in a series of steps: (1) Assume the location of the wing start; (2) find the shape of the mutation wing; (3) determine the total number of individuals, *n*(*x*), with an X chromosome of fitness *x* that the wing gives rise to; (4) from *n*(*x*) and *n*(*y*), determine the shape of the reassortment front by finding the establishment times for the front; (5) match the mutation wing to the reassortment front to fix the location of the wing start; (6) connect the mating rate λ/*L* to the nose fitness *Q* via the condition that the establishment times of new subpopulations at the front match the assumed mean speed.

The mutation wings for the reassortment steady state have the same dynamics as the wings for the asexual case of section 5. Let the X (*w* > 0) wing start have fitnesses 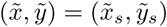 with 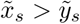 and define 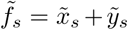. The front in the mutation wing is composed of lineages that began at the wing start and accumulated mutations on the X chromosome, as illustrated in fig. 3(a). Mutations are added at a rate of 1/τ_est_ = *f*/ *ℓ* so the X fitness increases at a speed *fs*/ *ℓ*. Therefore the relative front fitness, *f* = *X* + *Y* − *υt*, obeys eq. (17) or equivalently

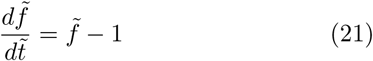

in rescaled units. After mutations accumulate for a rescaled time τ_*M*_ starting from 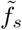 the relative fitness of this lineage is

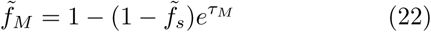

**Figure 3:**
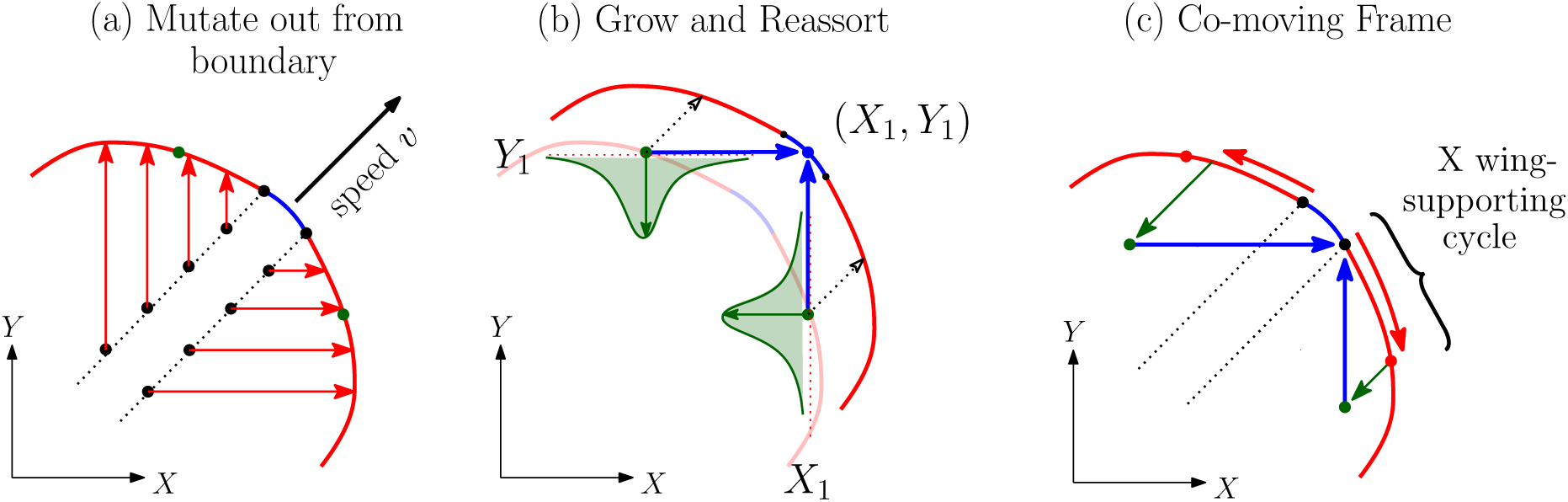
Schematic of the dynamics of the low mating rate steady state. The front is divided into the reassortment front (blue) and the mutation wings (red), which begin at the wing starts (black dot). (a) Lineages in the mutation wing started at a wing start some time in the past and accumulated mutations on only one of the chromosomes. (b) Subpopulations formerly at the front grow for a period of time and fall behind the advancing front. The conditional distribution of X-chromosome fitness, *X*_1_, is shown in green. The maximum of this distribution contributes the most copies of the X chromosome for reassortment (blue arrow). Reassortment results in establishment along the reassortment front and advances it. (c) The same dynamics shown in the moving frame. Mutating along one chromosome (red arrow) moves a lineage out along the wing. A subpopulation loses relative fitness while growing clonally (green arrow). After a period of growth, reassortment (blue arrow) can yield establishments at the front. The important X-wing-supporting cycle is shown. Note that the upper path shown that produces the Y chromosome for the X-wing-start, is not the Y-wing-supporting cycle, which is distinct (obtained by reflection of the X-wing cycle along the diagonal).

Because the mean advances, the relative *Y* fitness simply decreases to 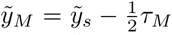 and the relative *X* fitness is found from 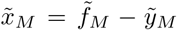. This determines the shape of the mutation wing 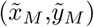 as a parametric function of τ_*M*_.

Various lineages in the mutation wings contribute to the total number, 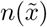, of chromosomes with fitness 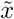 that are available for reassortment. As shown in fig. 3, these lineages mutated out from the wing start for different times before growing clonally. Consider a specific lineage that has mutated for a time τ_*M*_ from the wing start and then grows for a time τ_*G*_ without further mutations. During the growth, its relative *X* fitness decreases until it becomes 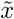. Thus τ_*G*_ is determined by

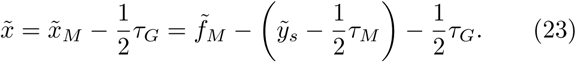

which gives τ_*G*_ as a function of τ_*M*_ and 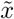. After this period of growth the size of this lineage is

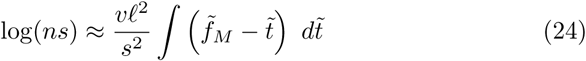

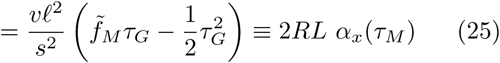

where *α* is a convenient rescaled quantity for log-populations and we have used eq. (20) to scale out *υ* in exchange for *R*.

The reassortment rate depends on the total number of individuals with an *X* chromosome with fitness 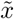 as shown in fig. 3. This total number is approximated by the integral

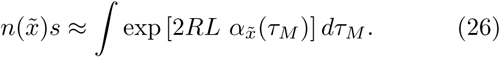

In the large *L* regime of interest this integral will be dominated by the largest subpopulation, so 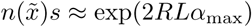 up to sub-exponential prefactors. The number of subpopulations contributing to the integral is roughly 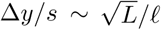. Even though this is large in the high speed regime (*L* ≫ *ℓ*^2^), the peak of 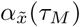 is narrow in the rescaled units, 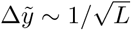.

The integral in eq. (26) is maximized when

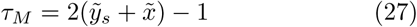

for which the fitness is 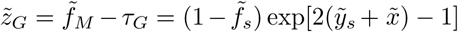 and the normalized log-population

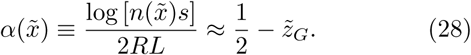

This determines how many chromosomes with fitness 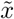 are available for reassortment.

We now find the shape of the reassortment front by using the condition for establishment by reassortment. From eq. (1), the rate at which mating produces individuals with 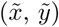 is

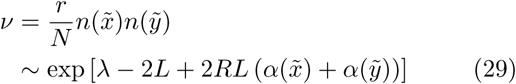

using the logarithmic measure for the mating rate, λ ≡ log(*Nr*). This rate *ν* grows exponentially with time at rate 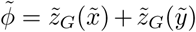, which is the sum of the fitnesses of the dominant subpops for 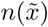 and 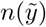. Because the growth rate of *ν* is not close to the fitness of the new individuals, i.e. 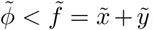, establishment is dominated by the first established individual, roughly when ∫^*t*^ *f ν e^ϕdt^* = 1 with solution *t* = − log(*νf/ϕ*) /*ϕ* ≈ − log(*ν*)/*ϕ*. This gives the establishment time for the front to advance by s from position 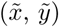:

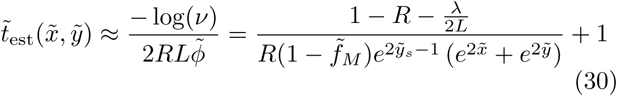

For a steady state, the whole front must advance at the same speed so the establishment time must be the same across the front. Since the 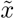 and 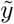 dependent part in eq. (30) must therefore be constant, the reassortment front has the simple form

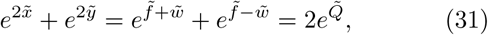

where 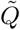 is the scaled nose fitness which occurs at 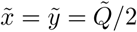. The position of the X-wing start, 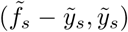, can now be determined in terms of 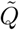 by matching the reassortment region (found from eq. (31)) and the mutation wing (inferred from eq. (22)) smoothly at the wing start. Matching derivatives at the wing start ensures that the mutation wing is “maximal,” i.e. mutants from nowhere else on the reassortment front could form a wing more advanced than the maximal one.

The nose fitness can now be obtained by requiring the establishment time to be *t*_est_ = *s/v* to match the steady state speed *υ*. In rescaled units, this is 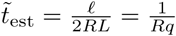 ≪ 1 and is thus much smaller than the terms on the right hand side of eq. (30) and can therefore be neglected: this is simply the continuous time approximation for the front dynamics valid for *q* ≫ 1. We can now substitute eq. (18) into eq. (30) to obtain

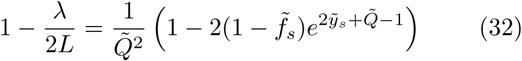

which, since we have already implicitly determined 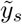 and 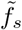 in terms of 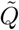, is an autonomous equation for 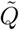 that can be solved numerically to give the shape of the fitness distribution. Note that the dependence on the mating rate enters only through the combination λ/*L*. The speed in un-rescaled units is then 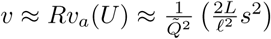, which is plotted in fig. 1.

The dependences on λ/*L* of several important quantities for the steady state solution are shown in fig. D.14. While the nose fitness 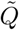 and the wing start fitness 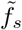 both decrease with λ/*L* because of the increasing speed, the fitness drop along the reassortment front, 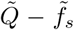, remains small throughout the low mating regime. But the width of the reassortment regime, measured by 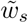, increases with the mating rate. The other important quantities characterize the X wing lineage that supports the X wing start by reassortment, as discussed more below. For increasing λ/*L*, the wing starts with a higher asymmetry in the fitnesses (higher 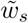) and smaller subpopulation sizes are needed for reassortment, so less time is needed for mutation (τ_*M*_) and growth (τ_*G*_).

The predicted speed increases linearly for small λ/*L*. But the inferred *υ* does not reach the expected sexual limit of *υ*/*υ_a_* (*U*) = 2 for high mating rates. Instead the solution approaches a cusp-like maximum value of *υ*/*υ_a_*(*U*) ≅ 1.56 at λ/*L* ≅ 0.84, which suggests that the *Ansatz* breaks down at or before this point. The breakdown occurs when the mutation wings in our *Ansatz* become unable to support each other via reassortment. The start of the Y-wing has fitnesses 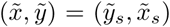 so it requires an X chromosome from the other wing with fitness 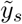. From eq. (27) we see that such a chromosome begins at the X-wing start and mutates out for a time 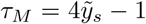. As the mating rate increases, the reassortment front becomes broader and 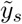 decreases, so less time is spent mutating. The *Ansatz* break down is when the 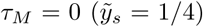, i.e. when no mutations are accumulated and the subpopulation that is crucial for mating grows up directly from the wing start. This corresponds to a mating rate of λ/*L* ≅ 0.81 (a mating rate lower than the non-sensical cusp-like maximum). For higher reassortment rates, a new steady state *Ansatz* is necessary: this must include additional reassortment events to support the mutation wings as discussed in the next section.

### 6.2. Fixation of mutations and genetic diversity

An important lesson from the steady state analysis is that cycles of mutation, growth, and reassortment are needed to support the advancing front. A particular cycle— the *wing-supporting cycle*—is most important because it is the path towards fixation for new mutations. Fig 3(c) shows this cycle: a lineage begins at the wing start, mutates, grows, and later feeds a new wing start for the same chromosome by reassortment. This new wing start then supports both the reassortment front and the next wing start. The λ/*L* dependence of the important quantities for this cycle— including the times for the important lineages to mutate and grow and the associated fitnesses — are plotted in fig. D.14. As the mating rate increases, the total (scaled) period of the wing-supporting cycle decreases roughly linearly from 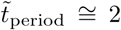 to 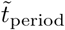 just below 1 when the low mating solution breaks down.

To understand the fixation process, it is useful to take a single mutation view of the evolution. We first review the behavior for asexual populations. The only mutations that have a chance of fixing arise in the low frequency, but very fit, subpopulations at the nose [28]. During the time that the nose advances by one mutational step, many mutations will establish and each mutant subpopulation subsequently grows. In order to continue to be competitive, a mutant lineage must be lucky enough to produce further mutations before its competitors. This process continues through a number—typically of order log(log *N*) [28]—cycles of single-mutation, establishment, and growth until the luckiest single lineage fixes in the nose. Soon after, the mutant lineage fixes in the whole population once the nose subpopulation has risen to dominate. It is this process that leads to fixations and generates the genetic diversity [28, 27].

The mutation fixing process in the two chromosome model is a generalization—albeit a subtle and complicated one—of that in asexual populations. Consider a chromosome with a mutation of interest that goes through the wing-supporting cycle. The chromosome will accumulate— up to order *q*—beneficial mutations and increase in copy number during growth. A few copies will establish at a later wing start due to reassortment. From there, these can accumulate additional beneficial mutations and continue through the wing-supporting cycle. New mutations that arise along the reassortment front or too far out in the wings are unable to fix because they occur on chromosomes that will not contribute to later reassortment. A mutation must first fix within the wing-supporting cycle in order to ultimately fix in the whole population. A mutant lineage must be luckier than competing lineages in establishing sooner at each of the many steps of the wing-supporting cycle. After a number of lucky rounds, a lucky mutation will have risen to be all but a tiny fraction of the population in the wing-supporting cycle and will soon fix.

In large populations, the important dynamics that maintain the mutation wings involve only low frequency sub-populations. Thus in experiments, the crucial dynamics underlying the evolution would not be seen unless the population were sequenced very deeply. We note, however, that in contrast to the asexual case, for very low reassortment rates some of the important dynamics takes place in large subpopulations: in this regime the sizes of the subpopulations that dominate reassortment to the wing-starts can be moderately large (albeit formally still exponentially small in λ).

### 6.3. Fitness distribution

The shape of the front determines the fitness distribution of the whole population. In the sexual limit (*r* → *s*) the fitness distribution is gaussian in both variables (indeed, it can be seen that the product of two steadily moving gaussians is formally a solution to the evolution equation, eq. (1) when mutations and fluctuations are ignored). But at low mating rates the distribution is far from gaussian especially in the low frequency regions important for the dynamics, e.g. the wing-supporting cycle in fig. 3. The shape of the full distribution depends on the front as

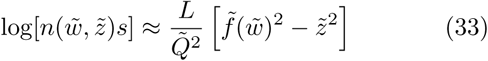

The distribution near its peak is controlled by the shape of the reassortment front near the nose, found in eq. (31):

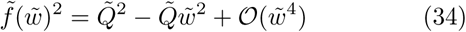

So the large frequency subpopulations near the mean are close to gaussian with an aspect ratio 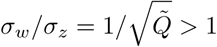. Non-gaussian corrections arise from the 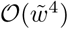 term and are noticeable when 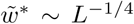, or 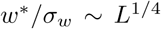. So for large *L*, the distribution appears gaussian for many standard deviations, but the important dynamics happen in the mutation wings where the distribution is far from gaussian.

The above asymptotic analysis yields an approximation to the steady state distribution, which becomes essentially symmetric near its peak as the mating rate decreases. But the asexual analysis of section 5 yields an anisotropic shape. This discrepancy is associated with subtle exchanges of limits of large *ℓ* and small λ. These are discussed briefly in Appendix Appendix C which addresses how the reassortment steady state approaches the correct asexual speed for finite *ℓ*.

### 6.4. Intermediate Mating Steady State

For λ/*L* ≳ 0.81—which we call “intermediate” mating although mating is still rare (*r* ≪ *s*)—a new *Ansatz* for the form of the steady state solution is needed because secondary reassortment events become important. In the low mating regime, larger λ means smaller subpopulations are needed for establishment by reassortment so smaller mutation wings—starting with smaller fitness, 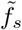, and further from the nose (larger 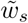)—can support the nose’s advance at faster speeds. For λ/*L* ≳ 0.81, however, the small X wing fails to produce subpopulations large enough to supply the X chromosomes needed for the Y-wing start. Instead, the Y wing is supported by reassortment from subpopulations that grew up from the reassortment front. The front for intermediate mating then becomes divided into three regions: the nose and wing regions described by equations in section 6.1, and new “re-mating” regions between the nose and wings that establish due to reassortment and also contribute to reassortment. Unlike for low mating, a chromosome can undergo two reassortment events (without accumulating new mutations in between) before losing too much fitness to reassort and establish at the front.

The secondary reassortment events make the dynamics more complicated but do not change the basic picture developed in the low mating case: a detailed discussion of intermediate mating is relegated to appendix Appendix B. One difficulty in the analysis is that the shape of the remating regions must be solved self-consistently since the Y re-mating region is a product of reassortment from the Y mutation wing and the X re-mating region (and likewise the X re-mating region depends on the Y re-mating region). In the appendix, we find the speed of the intermediate mating *Ansatz* without explicitly solving for the shape of the whole front. The results are plotted in fig. 1.

The intermediate mating *Ansatz* breaks down at λ/*L* ≅ 0. 99 when the two mutation wings cannot reassort to form the nose region. An additional multi-step process is needed for higher mating rates, and more and more complicated processes are likely needed as the sexual limit is approached, 1. e. λ → *L* or *r* → *s*.

## 7. Oscillations

Simulations of the two chromosome model reveal that oscillations are an important feature of the dynamics, especially for large populations. Park and Krug [37] noted the presence of oscillations but did not study them beyond describing the dynamics of the fitness distribution as a “breathing traveling wave.” As shown in fig. 4, the oscillations involve a cycle of diversity buildup through mutation followed by purging of much of the diversity by reassortment and subsequent selection. We will show that the dynamics of the oscillations are related to the dynamics within the steady state solutions discussed above. The oscillation dynamics differ for low and intermediate mating rates, but here we focus solely on the low mating oscillations, which capture the important features of the dynamics, and discuss the intermediate case in appendix Appendix B. The oscillations are clearly observable for the stochastic dynamics and become more regular for *q* ≡ 2*L*/ *ℓ* ≫ 1 large, see fig. 4. Thus the deterministic simulations (described in section 4) are very useful for characterizing and understanding the oscillations.

**Figure 4:**
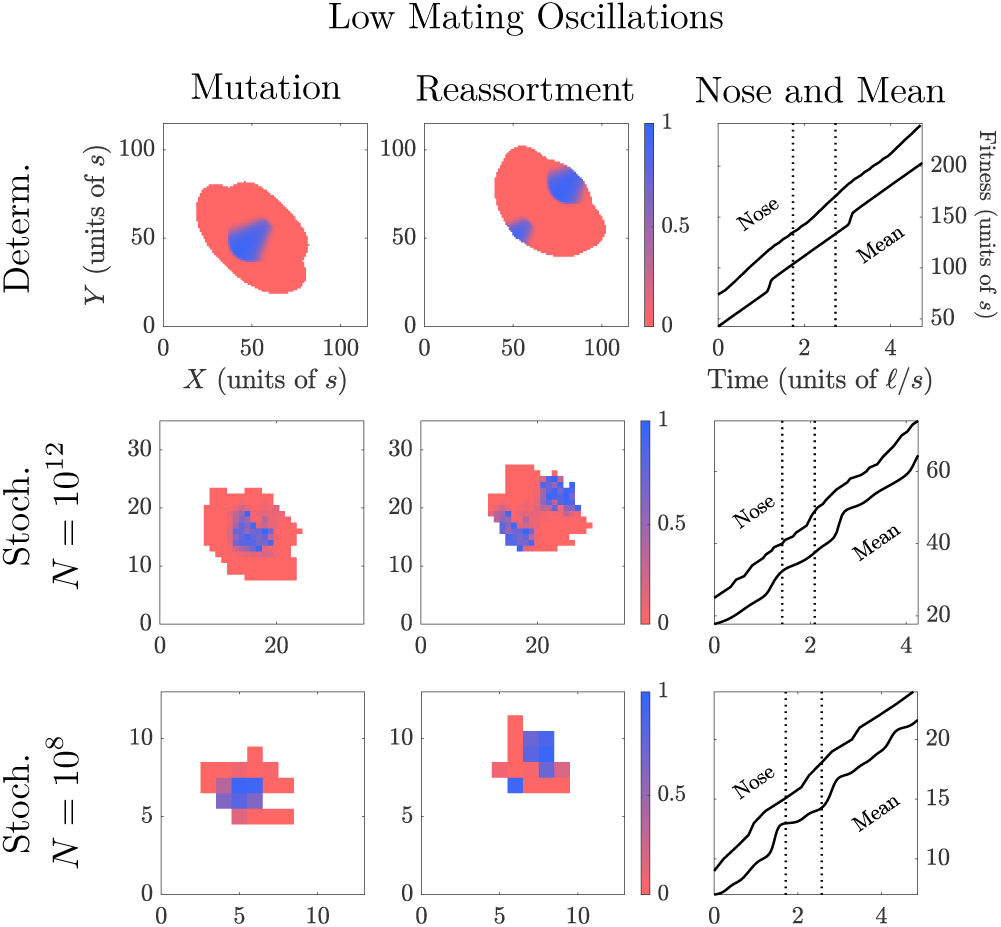
Oscillation cycles illustrating the fitness distribution during the mutation phase (left column) and the reassortment phase (middle column) that occur in each cycle. Top row: deterministic approximation to the dynamics; lower rows: stochastic dynamics. All the non-zero subpopulations are shown with their color indicating the fraction that was established by reassortment (bluer) or mutation (redder). The oscillation dynamics are shown at low mating rates: λ/*L* ≈ 0.5. The righthand column shows the speed of the nose and mean, dashed lines corresponding to the times of the snapshots shown. The nose speed increases during the reassortment phase and, through exponential growth of the prior nose populations, the effects of this are sharpened into a jump in the mean fitness roughly a time i/s later. The stochastic simulations have *N* = 10^12^, *s* = 10^−2^, 2*U* = 10^−4^, *r* = 10^−7^ (*q* ≈ 9, *ℓ* ≅ 5.3) and *N* = 10^8^, *s* = 0.03, 2*U* = 10^−6^, *r* = 1.8 × 10^−5^ (*q* ≈ 2.7, *ℓ* ≅ 11). These agree qualitatively and semi-quantitively with the deterministic simulations, which are valid in the continuous limit that obtains when *q* ≡ 2*L*/ *ℓ*—which determines the width of the fitness distribution—is large.

The oscillation dynamics for the deterministic and stochastic simulations can be seen in the patterns of establishment shown in fig. 4. Each fitness class of the current population distribution is colored according to whether its establishment was dominated by reassortment or by mutation. At any point in the cycle and at each position along the front, either reassortment or mutation strongly dominates as occurred for the different sections of the front in the steady state *Ansatz*. The oscillation cycle can thus be cleanly divided into two phases: a reassortment driven phase of establishments at the nose that will eventually advance the whole population, and a mutation driven phase that both advances the nose and creates the diversity for reassortment to later act on.

The oscillation dynamics over one period resembles the cycles of mutation, growth, and reassortment described in the steady state analysis. During the reassortment phase, reassortment advances the central part of the front. The front advances faster than the mean, so the nose (the fittest part of the front) becomes more and more fit (relative to the mean) during this phase. These fitter subpopulations from the nose grow especially fast until eventually one takes over the bulk of the population, causing the mean fitness to jump sharply. Almost immediately after the mean jumps, the population has too few chromosomes with high relative fitness that could possibly reassort to the front, thereby quickly ending the reassortment phase. But the front can still advance by mutation.

During the subsequent mutation phase, the central region of the front advances more slowly, as shown in the righthand column of fig. 4. But more important for the future are lineages in the wings that accumulate mutations asymmetrically—some predominantly on one chromosome, some predominantly on the other—extending both the wings. The growth of these wing subpopulations eventually creates enough high fitness chromosomes that reassortment can establish new subpopulations at the front: this starts the reassortment phase again and the speed of the nose increases. In summary, over each period, reassortment rapidly advances the nose which causes the mean to jump forward and stops reassortment to the front. Then mutation and growth in the wings produce the higher fitness chromosomes that are again able to reassort to the front and advance the nose.

Looking in more detail at the nature of the oscillations, it is apparent from the figures that the variations in the speed of the mean are much larger than those in the nose speed: indeed, a sudden increase in the speed of the nose becomes sharpened into a jump in the mean fitness because of the exponential growth of competing subpopulations. We illustrate these dynamics through a simple toy example. Assume that the nose and the mean have been traveling at speed *υ*_1_ for some time and at *t* = 0 the nose speed increases to *υ*_2_. (This scenario is a surprisingly good approximation for the low mating oscillations, as seen in fig. 4). During the mutation phase, the nose and mean settle into traveling at roughly the asexual speed until the reassortment alters the nose speed. The mean is unaffected by the change in nose speed until the new subpopulations reach a size comparable to N. New subpopulations establishing at time *t_e_* > 0 start with a larger relative fitness, *f*(*t_e_*) = *Q* + (*υ*_2_ − *υ*_1_)*t_e_*. Here *Q* is the original nose fitness, which should satisfy *Q*^2^ = 2*υ*_1_*L* from eq. (18) if the nose and mean have travelled at υ_1_ for longer than τ_nm_ = *Q*/*υ*_1_ which we will assume. A subpopulation will reach a size ~ *N* at a time *t_N_*(*t_e_*) when

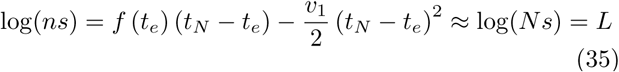

When the first subpopulation reaches a size *N*, the mean will jump to the (absolute) fitness of that subpopulation. This subpopulation established at the time 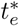 that yields the earliest time 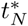, and hence satisfies *dt_N_*/*dt_e_* = 0. The jump in mean fitness is simply the relative fitness of this subpopulation immediately before the jump:

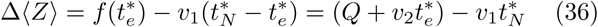

The extremal condition equates the jump size to Δ〈*Z*〉 = (*υ*_2_ − *υ*_1_) 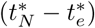. The detailed solution can be expressed in terms of the speed ratio *β* = *υ*_2_/*υ*_1_:

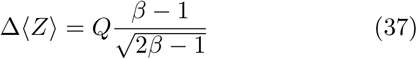

Thus with *β*−1 not small, the jump size is some substantial fraction of *Q* as seen in fig. 5. It is interesting to note that the largest jump sizes—roughly 0.5Q—correspond to *υ*_2_ ≈ 2*υ*_1_ which would be expected if the wings mutated out at the same speed *υ*_1_ as the nose advances by mutations.

**Figure 5:**
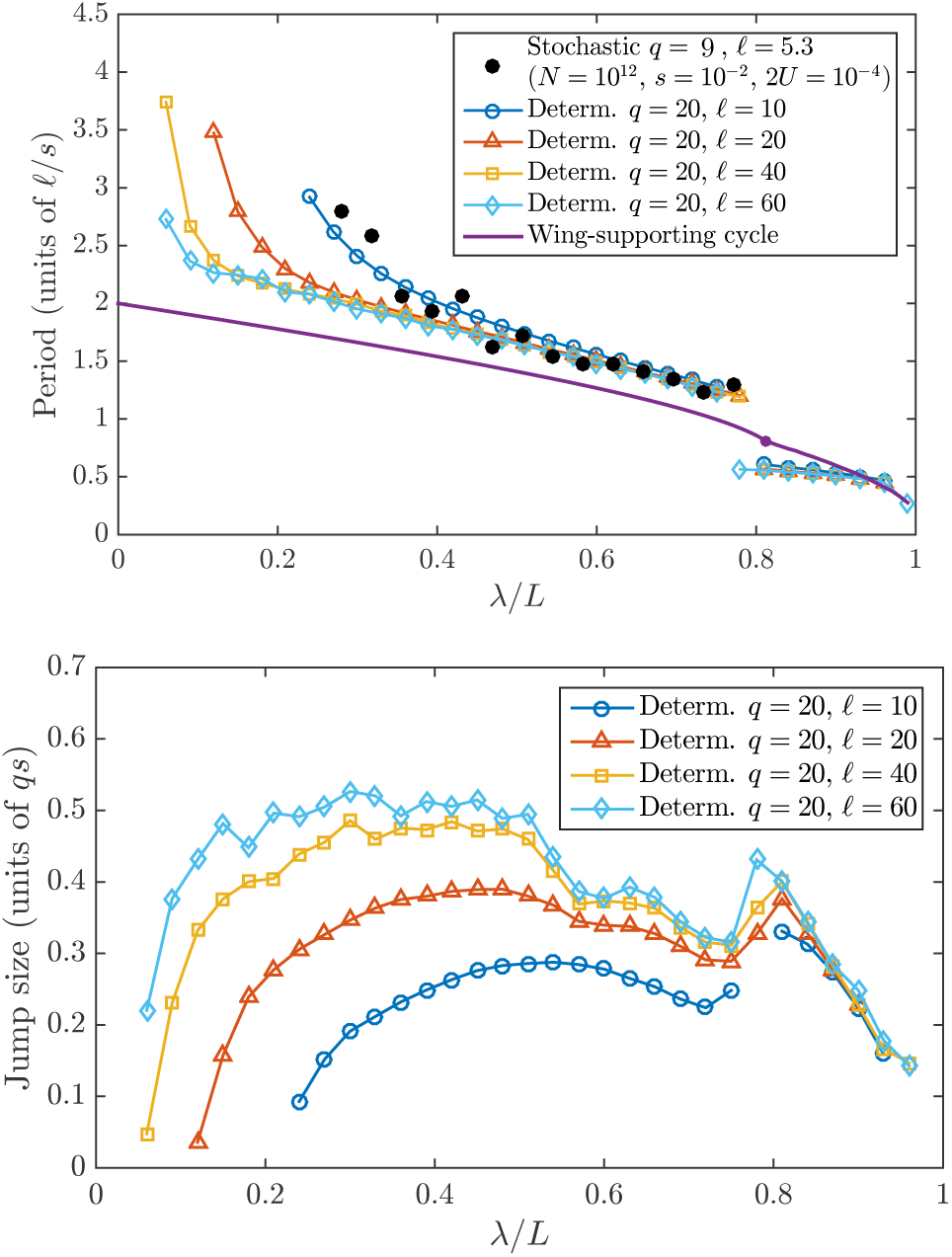
Quantitative properties of the oscillations: period (top) and jump size of the mean fitness (bottom) in the deterministic and stochastic (black dots) simulations. For the stochastic simulations the period was extracted from the power spectral density for five simulation runs of 300 *ℓ*/*s* time-steps. For comparison, the period of the wing supporting cycle of the steady-state is shown (purple line). The deterministic simulations show that the period diverges at low λ/*L* as the dynamics approach the asexual limit. The period can become several times greater than the asexual nose-to-mean time, *ℓ*/*s*, because the distribution must relax nearly to the asexual steady state before the next reassortment phase begins. At λ/*L* ≅ 0.8 there is a period-halving bifurcation corresponding to the transition to the intermediate mating regime of the steady state.

The dependence on λ/*L* of the important oscillation characteristics—the period and the magnitude of the jump of the mean fitness— are plotted in fig. 5. The oscillation period from the deterministic simulations is seen to be similar to the period of the asynchronous wing-supporting cycles of the steady state for large *ℓ* Furthermore, there is a period-halving bifurcation near λ/*L* ≈ 0.8 due to a change in the oscillation dynamics. This bifurcation occurs close to the transition from low to intermediate mating for the steady state solutions. Together these suggest a strong connection between the oscillatory and steady state dynamics. This can be understood in terms of the lineages that are most important for reassortment. In the oscillatory dynamics, the mutations wings trace back to a small segment of the front—not necessarily near the nose—at the end of the previous reassortment phase, shown schematically in fig. 6. This segment of the front is analogous to the steady state wing start that produces the full mutation wing and the dynamics that support the key parts of the front are the same as the wing-supporting cycle. The main difference is that speed of the mean, *υ*(*t*), which influences growth and establishment rates, is not constant and instead depends on the past nose fitness. In the steady state solution, many wing-supporting cycles take place in parallel but out of phase with each other, each depending on when their wing start establishes. For the oscillations, the uniform distribution of phases of the cycles breaks down and the cycles lock together, most likely due to instability from the delayed feedback between the speedup of the nose and jump in the mean. Although we have not attempted a full linear stability analysis of the steady state, deterministic simulations of a simplified caricature shows that its steady state is indeed unstable to oscillations. This caricature has establishment by reassortment only right at the nose but that is enough to incorporate the delayed feedback structure.

**Figure 6:**
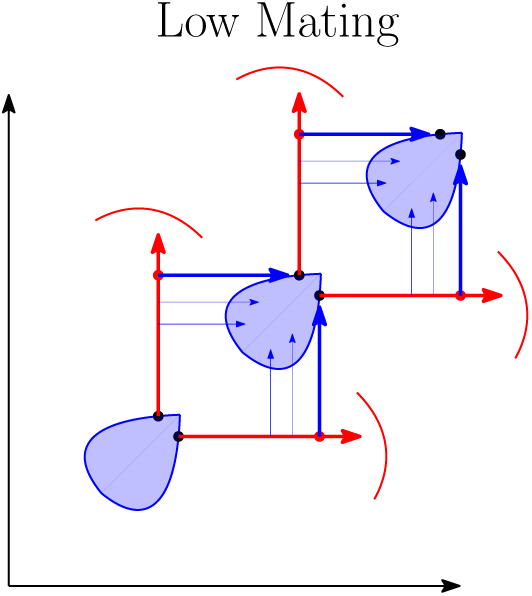
Diagram of the oscillations at low mating. The blue convex regions represent subpopulations that established by reassortment. The oscillation dynamics resemble the steady state cycles in fig. 3 that support the wing start: a lineage mutates (red arrow) predominantly on a single chromosome before reassorting (blue arrow). The oscillations appear as a single dominant cycle, as opposed to the many overlapping cycles in the steady state.

At small λ/*L*, the period depends on *ℓ* but appears to converge to a limit for large *ℓ* that is close to the period of the wing-supporting cycle in the steady state. However the jump size depends strongly on *ℓ*. This suggests that the *ℓ* dependence of the speed for low mating in fig. 1 (discussed more in Appendix Appendix C) is largely due to the different jump sizes.

### 7.1. Oscillations for *ℓ*x2192; ∞

A quantitive description of the deterministic oscillations for general *ℓ* and λ is difficult because of the complicated feedback structure of the dynamics. The mean fitness depends on the nose fitness at previous times and the nose fitness depends on the time for the wing lineages to mutate, grow clonally, and reassort which all depend on the history of the mean fitness. The mutation rate for the wings also depends on the local shape of the front because of the difference between two-parent and one-parent mutation (described in section 5) which is controlled by *ℓ*. However, the essential features of the oscillations can be captured by a much easier to understand limit: *ℓ* → ∞ and λ → 0 in the continuum limit of large *q*. For large *ℓ*, the difference between two-parent and one-parent mutation rates is negligible and the two-chromosome and one-chromosome speeds are the same: *υ_a_*(2*U*) = *υ_a_*(*U*) = (2*L*/ *ℓ*^2^)*s*^2^. For λ/*L* = 0 a subpopulation contributes to reassortment simply when its fitness equals the mean fitness and its size is roughly N. As we will show, the oscillations result in a sizable speedup even though λ/*L* → 0—a surprising result.

Let us consider what happens for *r* ~ 1/*N*—i.e. only 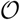(1) matings per generation in the whole population. An initially compact fitness distribution evolves under asexual dynamics with a speed that approaches the asexual speed. The middle part of the asexual front (which for large finite *ℓ* has the form 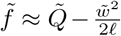 as derived in Appendix A), becomes, for *ℓ* = ∞, *flat* with well-defined ends. This front later gives rise to a *ridge* at the mean fitness of width *W_R_*, along which the subpopuations have the same size: each of order *N*. At some point the ridge becomes wide enough that reassortment can lead to establishment at the nose—the center of the flat front. For *r* ~ 1/*N*, only subpopulations located on the ridge (i.e. with 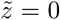) can feed the nose, thus with a nose at 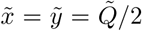, the sub-populations feeding it must be at 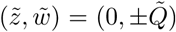: these will feed the nose at a rate of order one. The reassortment will then advance the front faster than the asexual speed until the mean jumps, which quickly ends the reassortment phase.

Because of the flat front and ridge, the geometry of the dynamics is much simplified. The full oscillation dynamics reduce to a one-dimensional description: knowledge of the nose fitness *F*(*t*) for all previous times *t′* < *t* is enough to determine the future dynamics. It is convenient to start from the time, *T*_0_, at which the mean jumps: call the nose fitness at that time, *F*_0_. The first plot in fig. 7 shows how mutation from this last point of reassortment—denoted by a blue dot—results in an expanding triangular region with a flat front. Subpopulations growing from the flat front established at the same time and therefore have the same population size. They form a ridge when they reach maximum size—each 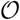(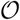)—and their fitness equals the mean fitness 〈*Z*(*t*)〉.

**Figure 7:**
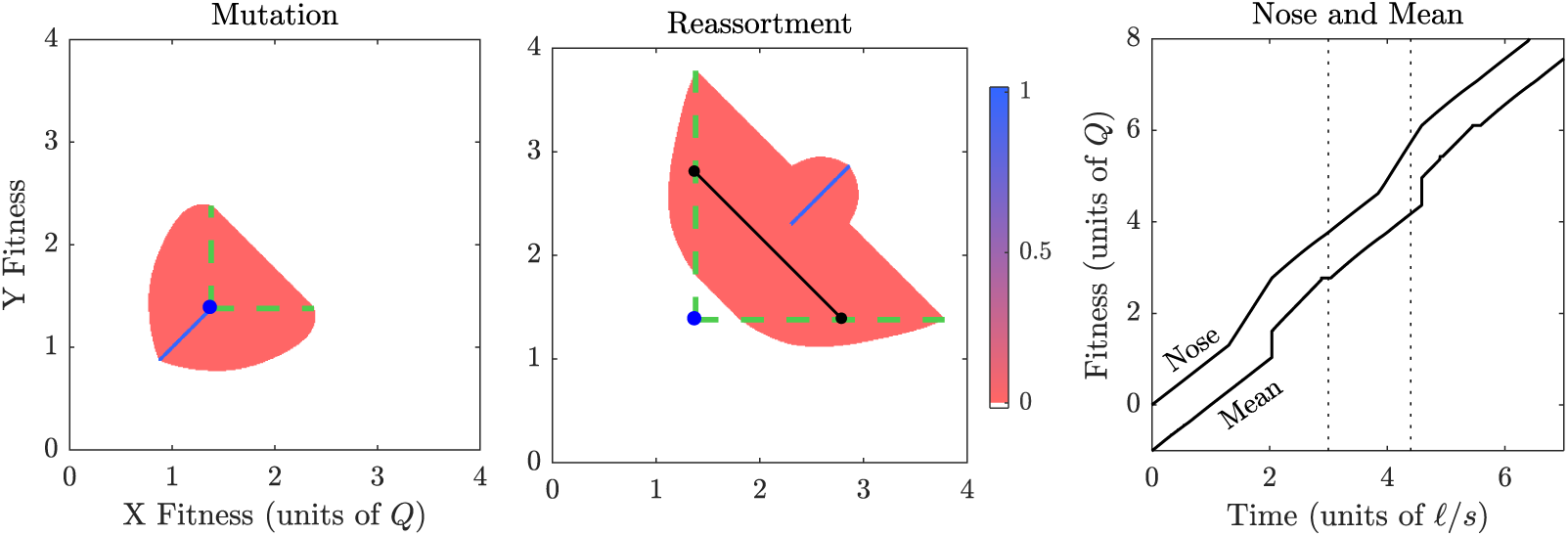
The oscillation dynamics for *ℓ* → ∞ and λ → 0 have a simple geometry. During the mutation phase, left, a flat front mutates out from the last reassortment point—indicated by the blue dot—forming an expanding triangular region bounded by the dashed green lines. The front is flat because the difference in establishment times between one parent subpopulation mutating and two parent subpopulations is negligible in the *ℓ* → ∞ limit. A set of subpopulations each reach a size log(*ns*) = *L* needed to reassort, center, when their fitness equals the mean fitness: these form a high population ridge indicated by the black line. The X-most and Y-most edges—indicated by small black dots—then advance the nose via reassortment. Despite the simple geometry, the feedback between the nose and the mean leads to a nontrivial trajectory for the nose and mean fitnesses (right hand figure). The plots are from deterministic simulations of the *ℓ* → ∞ and λ → 0 dynamics described in the text.

The highest fitness point to which reassortment can occur is when the reassorters are at the opposite ends of the ridge. The X-most and Y-most edges have 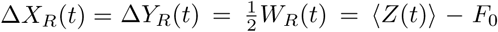 where the last equality follows from the geometry of the triangular front that gave rise to the ridge populations. Reassortment results in establishment of a subpopulation with fitness 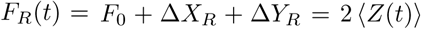 that will advance the front if this is greater than the current front fitness. Let the time at which this reassortment starts be T_R_. During the subsequent reassortment phase, the nose fitness is *F_R_*(*t*) which advances at twice the speed of the mean, 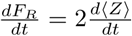. Although the nose advances due to reassortment, other new subpopulations—even those immediately away from the nose—establish due to mutation. This is because lineages descending from previous nose subpopulations are especially fit and can advance the front by mutation quicker than reassortment except right at the new nose. Thus the reassortment region in fig. 7 is limited to the thin blue line shown.

The delayed advance of the mean can generally be determined from the nose fitness *F*(*t*) by considering the time for nose subpopulations to reach size *N* (as in our illustrative analysis of jump sizes in eq. (36)) A subpopulation that established at time τ has a size at time *t* of

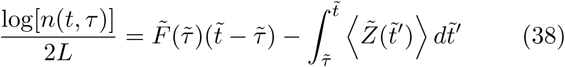

The rescaled mean fitness is then

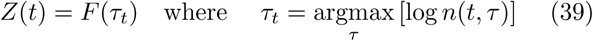

or simply the fitness of the subpopulation with largest size.

During the mutation phase starting at *F*(*T*_0_) = *F*_0_, the speed of the nose is 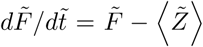. When combined with the previous results for *F_R_*(*T*) = 2〈*Z*(*T*)〉 − *F*_0_, the nose fitness is

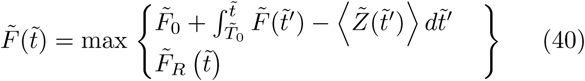

The reassortment phase begins when the max switches to *F_R_* at time *T_R_*. The faster nose speed during reassortment will later result in a jump in the mean fitness at time *T_J_*. This happens when the τ_*t*_ jumps discontinuously from a value τ_*t*_ < *T_R_* to a time τ_*t*_ > *T_R_* which means that a subpopulation established during the reassortment phase has taken over as the largest subpopulation. At time *T_J_* reassortment stops and a new mutation phase begins: between *T*_0_ and *T_J_* the front has thus completed a full cycle. If this cycle reaches a periodic steady state with jumps at regular intervals, the system has stable oscillations: this is indeed what we find in direct simulations of this one-dimensional non-linear delay-dynamical system.

The trajectories of the nose and mean fitness in the infinite *ℓ* limit analyzed above are shown in fig. 7. Surprisingly, the oscillations result in a speed of *υ* ≅ 1.3 *υ_a_*(2*U*) implying that the infinite *ℓ* speed jumps discontinuously at λ/*L* = 0! This is a consequence of the reassortment phase lasting for a substantial fraction of the period of the oscillations. Thus the interplay between mutation and reassortment even for mating rates as low as *r* = 1/*N* can yield ≈ 30% of the total possible benefit of mating. Note that, formally, as long as *NU* is very large, even in the large *ℓ* limit, *r* ~ 1/*N* corresponds to *r*/*U* ~ 1/(*NU*) ≫ 1 and thus reassortment rate much less than beneficial mutation rate.

Since we found that 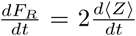 in the reassortment phase, we can use our previous calculations in eq. (37) for the dependence of the size of the mean jump on *β*, the ratio of the nose speed before and after reassortment starts, to estimate the jump size. With *β* ≈ (2*υ_a_*)/*υ_a_*, this gives 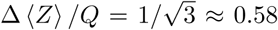, roughly what is observed. Although the infinite *ℓ* limit might seem pathological, the jump sizes in fig. 5 saturate to a similar value for large *ℓ* because of the same underlying dynamics. The X-most point that can produce reassorters into the nose advances in the X direction at roughly the mean speed, and similarly for the Y-most point. So the nose advances due to reassortment at roughly twice the mean speed. A discussion of the general dependence of the speed and jump size on *ℓ* can be found in Appendix Appendix C. The behavior depends on how close the population at the end of the mutation phase is to the asexual steady state versus the infinite *ℓ* limit: the latter having large oscillations and being far from the steady state.

## 8. Fluctuations

The deterministic approximation gives a good qualitative and semi-quantitative picture of the dynamics, but leaves out the fluctuations that are visible in the stochastic plots of fig. 4. While we have not analyzed these in detail, we outline here some general qualitative and quantitative features of the fluctuations.

In the asexual case, fluctuations in the speed of the nose give rise to jumps in mean fitness. Small asexual nose fluctuations are of order 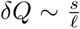 and are only correlated for a short time, roughly a single establishment [40]. The fluctuations get amplified by exponential growth and produce jumps in the mean with an exponential distribution of sizes with mean of order 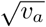, which scales with *Q* as 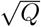. The nose-driven stochastic jumps of the mean stabilize the system against large oscillations. The nose is unstable on short times because an early establishment induces an even earlier establishment next. This leads to an accelerating nose speed that can only be corrected by feedback a time τ_nm_ later. However, the larger resulting jump in the mean (due to the earlier nose fluctuation) will dominate the future nose dynamics and prevent the nose from running away [40]. In contrast to the asexual case, for sexual dynamics the oscillations involve large changes in nose speed that are sustained for a long time, resulting in large jumps in the mean of order *Q*. Thus the asexual jumps and oscillation jumps can be distinguished by their scaling with *Q*.

Asexual mutational fluctuations modify the behavior of the two chromosome asexual steady state of section 5. The shape of the two-dimensional front we derived does not strictly correspond to a steady state because the stochastic dynamics allows the average w coordinate to diffuse over time. A part of the front will fluctuate ahead and descendants of this part grow faster and mutate out to form a new front, shifted relative to the previous one. Our steady state then roughly represents the “typical” shape of the distribution when this lateral diffusion is suppressed or when large fluctuations have not occurred in the recent past. (Note that even in the one chromosome asexual model, the amplified effects of the nose fluctuations make interpretation of the steady state distribution as an ‘‘average” already subtle: it loosely represents a “median” shape in the frame of the mean fitness.)

Reassortment adds another source of stochasticity and a different feedback structure. Nose fluctuations now depend on the stochasticity of two separate lineages undergoing a cycle of mutation, growth, and reassortment. For the oscillations, the start time of the reassortment phase is stochastic and this will later have a large effect on the time and magnitude of the mean fitness jump. Establishments due to reassortment are more stochastic than those due to mutation. For reassortment the first established lineage is likely to dominate, but for mutation many independent secondary lineages will also contribute substantially and their combined effect is to decrease the stochasticity. [28, 40] Thus it is not surprising that the added stochasticity from reassortment results in greater diffusion of the mean and nose than in the asexual case, as found in fig. 8.

**Figure 8:**
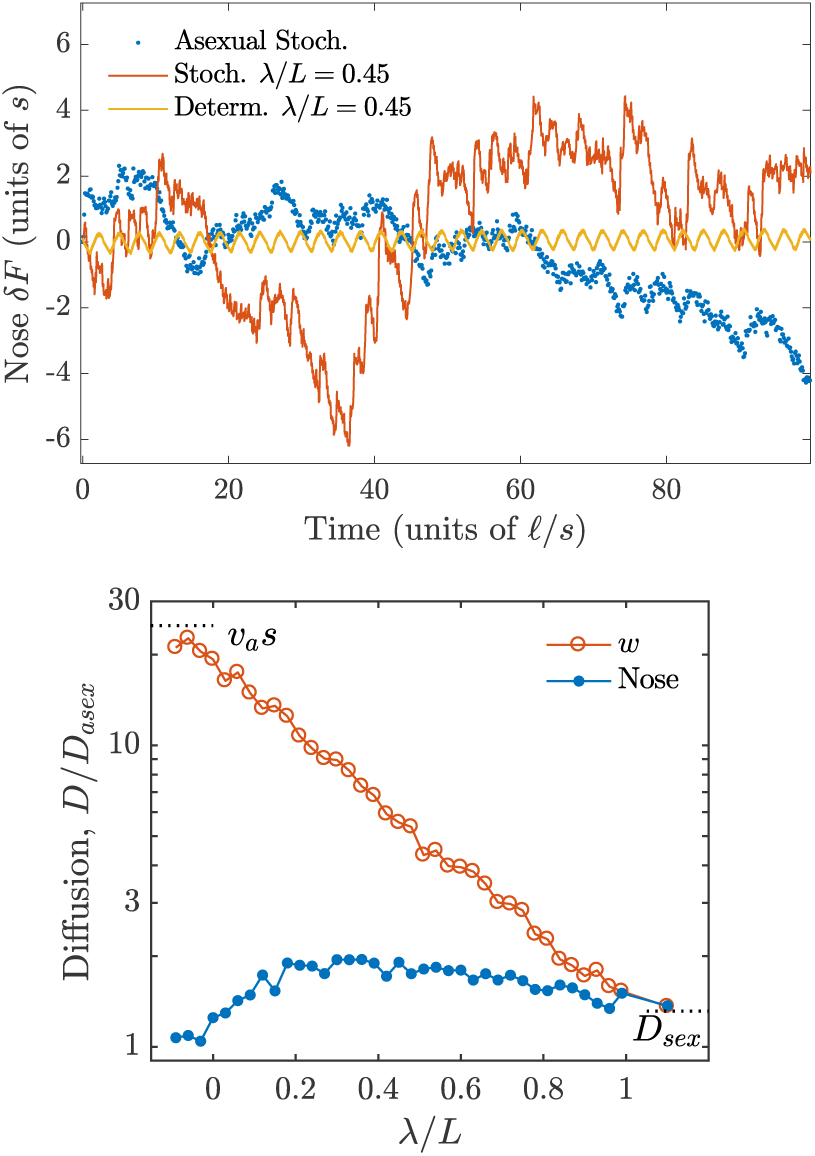
(a) Fluctuations in the nose fitness *δF* with reassortment (red) with λ/*L* = 0.5 and without reassortment (blue). A deterministic simulation (yellow) shows the period of oscillation, which can also be seen in the stochastic simulations. (b) Diffusion constants for the nose fitness and the mean transverse fitness 〈*W*〉 = 〈*X* − *Y*〉 plotted on a log scale. The results are normalized by the theoretical prediction for the asexual case, *D_asex_* = 2π^2^(*s*/ *ℓ*_2_)^3^/3 with *ℓ*_2_ ≡ log(*s*/2*U*). The other limits indicated, *D_w_* = *υ_a_s* and *D_sex_* = 2*D_asex_*(*ℓ*_2_/ *ℓ*)^3^, are explained in the text. For the nose, diffusion is greatest for reassortment in-between the asexual and sexual limits when the jumps in mean fitness due to oscillations are greatest. Parameter values for both plots: *N* = 10^12^, *s* = 10^−2^, 2*U* = 10^−4^ (*q* ≈ 9, *ℓ* ≅ 5.3). Diffusion constants were calculated from 10 simulation runs of length 500 *ℓ*/*s*.

Figure 8 shows that diffusion is greatest for mating rates when the dynamics are neither asexual or fully sexual. In the asexual limit, the diffusion constant is predicted to be *D_asex_* = 2π^2^(*s*/ *ℓ*_2_)^3^/3 with *ℓ*_2_ ≡ log(*s*/2*U*) [40]. The fully sexual limit is simply related to the asexual limit as the two chromosomes become completely unlinked and each evolve asexually (with *ℓ* instead of *ℓ*_2_) so the diffusion of the nose is twice the diffusion of a single chromosome, *D_sex_* = 2*D_asex_*(*ℓ*_2_/ *ℓ*)^3^, where the (*ℓ*_2_/ *ℓ*) factors convert *D_asex_* to the single chromosome result. The sexual limit of the transverse diffusion of *w* in this limit is also *D_sex_* since the sum and difference of two unlinked fitnesses have the same diffusion. Park and Krug [37] derived the asexual limit of the transverse diffusion to be simply *D_w_* = *υ_a_s* using the fact that mutations fix at a rate *s*/*υ_a_* and occur randomly on one chromosome or the other. They showed that this initially high diffusion decays rapidly with reassortment. Figure 8 shows that this decay is surprisingly exponential over the full range of λ values: we have not investigated the source of this behavior.

Near the transition between low and intermediate mating rates (at λ/*L* ≈ 0.8), in the deterministic approximation there is a small window of bi-stability between the low mating oscillation style and the intermediate one. Fluctuations can lead to transitions between the two different oscillation modes. The transitioning fluctuations become rarer for larger *q* so a longer simulation time is needed to average over the two oscillation modes. In the deterministic simulations (roughly the *q* → ∞ limit), the bi-stability manifests as hysteresis when the mating rate is slowly varied: the current oscillation mode depends on the past mating rate and whether the mating rate has increased or decreased to the current value. In fig. 8 the bi-stability does not result in greater diffusion around λ/*L* ≈ 0.8 because the parameter values used are stochastic enough to smooth over effects due to the transitions between modes. This suggests that bi-stability is unlikely to be important for realistic population sizes.

## 9. Discussion

When sex is very rare, mating has negligible effects on already established subpopulations which simply increase in size due to clonal growth. But even with additive effects of mutations, mating is crucial for the creation of novel, very fit genetic combinations that will drive the future evolution. Offspring fitness after mating depends on the genetic relatedness of the parents, so in the rare sex regime it is crucial to track the details of the genetic diversity in the population. In large populations of size *N*, the important aspects of this diversity involve subpopulations with very anomalous past mutational and mating histories. The crucial properties of the diversity and how it is determined in an evolving population cannot be captured by a few statistical properties. The simplifying feature of the two chromosome model with reassortment is that the important relatedness and dynamics are fully captured by tracking only a two dimensional fitness distribution. For this model, we have shown explicitly how the properties of rare subpopulations with anomalous history control the dynamics. And these dynamics are complex, involving long cycles of mutation, clonal growth, and mating to produce the unusually high fitness individuals whose descendants will dominate the future evolution. We find that the speed of evolution depends logarithmically on the mating rate, *r*, so that sizable speedups can occur for very small *r*, and that the ratio of log(*rN*) to log *N* is the important combination of parameters. While the detailed dynamics of evolution of large populations with very low rates of mating or lateral gene transfer will surely be different than the two chromosome model, we expect that such logarithmic dependence on the recombination rate as well as the dominance of the dynamics by subpopulations with very anomalous histories will be rather general. These features, as well as how general the cyclical dynamics might be, we discuss below.

### 9.1. Summary of qualitative picture of two-chromosome model

For rare mating in the two chromosome model, producing fitter offspring than mutation can produce requires a large population of anomalously high fitness chromosomes. Large subpopulations are produced by clonal growth from populations that established at the front — the high fitness edge of the two-dimensional fitness distribution. Our steady state analysis shows that the high fitness chromosomes that can advance the front via reassortment arise in the “wings” of the front which are produced by anomalous lineages that mutate predominantly on a single chromosome. These anomalous subpopulations then grow clonally to sizes sufficiently large for significant mating. The predominant matings involve parents with one very fit chromosome and one average fitness chromosome. By analyzing how the wings are produced by a cycle of mutations, growth, and mating, we find the steady state speed of evolution. The “wing cycle” dynamics driving the evolution involves only low frequency subpopulations. This is a generalization of the mutational dynamics at the nose of asexual populations which controls the future evolution (as well as dominating the fluctuations and diversity statistics) [44, 28]. A key difference, in addition to the two-dimensionality of the fitness distribution, is that the wing cycle is much longer than the times between mutational steps: it takes of order the time for subpopulations to reach a size ~ *N*.

An unexpected feature of the two chromosome dynamics is sustained oscillations (first observed in the simulations of Park and Krug [37]), which resemble the wing cycle of mutation on a single chromosome, clonal growth, and reassortment. We find that the oscillations cause the evolution of the whole population to be separated into distinct mutation-driven and reassortment-driven phases. Although the scalings of both periods with log(*Nr*) are quantitatively similar, the oscillations speed up the evolution because they result in large jumps in the mean fitness by a substantial fraction of the difference, *Q*, between the mean and maximum fitness of the population. Counterintuitively, rare reassortment results in periods of reduced diversity: after reassortment creates a set of fitter subpopulations, selection purges less fit subpopulations more rapidly. And this stronger selection later results in a jump in the mean fitness. Since a small subset of lineages contribute to reassortment each cycle, the oscillations act as a bottleneck and are thus important for understanding the diversity which we have not analyzed. The stochasticity of the reassortments gives rise to stochasticity in the advance of the mean fitness which is much larger than that for asexual populations, as seen in fig. 8.

In the rare sex regime, the speed of evolution increases over a broad range of mating rates from the inverse population size, 1/*N*, up to the selective strength of mutations, *s*. Dependence on the mating rate comes through the parameter combination log(*Nr*)/log(*Ns*), which varies from zero to one over this range. Since the dynamics of course depend on the total population mating rate, *Nr*, this ratio of logs is only significant because it also accounts for the overall N dependence of the asexual to sexual transition. For very large populations, log(*Ns*) sets the overall fitness scale such that the mutation and growth dynamics can be rescaled to have no *N* dependence. This was shown explicitly for the steady state solution but holds in general, including for the oscillations. The exponential growth of subpopulations implies that the natural rescaling of subpopulation size is *α* ≡ log(*ns*)/log(*Ns*) which is typically of order one in the interior of the population distribution and ranges from zero for a single individual to one for the largest subpopulation with size ~ *N*. Reassortment from individuals in different parts of the fitness distribution will establish a new subpopulation *n*(*x, y*) roughly when the influx rate due to reassortment, *r n*(*x*)*n*(*y*)/*N*, is of order one—with *n*(*x*) as the total population with X-chromosome fitness, *x*. The condition for establishment by reassortment is then simply 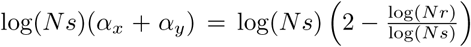showing that log(*rN*)/log(*Ns*) is the natural combination that determines the quantitative effects of reassortment.

### 9.2. Fixations and diversity

Even though mating is continually happening, the oscillation dynamics make it appear as though the population were undergoing periodic mating—except that the period is determined by the other evolutionary processes instead of being fixed externally. The period of oscillation sets a natural fixation timescale because the population in the next cycle descends from only a small number of subpopulations that successfully mated to the front. This has important implications for beneficial and neutral diversity statistics and the structure of the genealogies as many small mutations accumulated in the mutation phase can fix at once. But in order for a mutation to fix, it must arise in the right part of the front and then its lineage must be anomalously lucky in accumulating further mutations, then lucky in reassorting to the front, and then lucky in continuing through several such cycles, in order to outcompete other lineages that arose around the same time.

Our analysis of the deterministic steady state approximation to the dynamics gives qualitative and some quantitative hints at how mutant lineages arise, fluctuate, and some lucky ones eventually fix. However the interplay between the oscillations and fluctuations make it difficult to analyze the coalescent and diversity statistics that this process gives rise to. Analyzing these for the two chromosome model should be a fruitful direction for future work. We expect that the coalescent process will not be in the same universal class as either the conventional neutral Kingman coalescent or the Bolthausen-Sznitman coalescent predicted for large continually evolving asexual populations [27, 28], although, we do expect that, like the latter, the phylogenies will be characterized by multiple mergers.

### 9.3. Quantitative results: beyond asymptopia

Our primary results are valid in asymptotic regime in which the logarithmic parameter *L* = log(*Ns*) is large. Further simplifications occur when *ℓ* = log(*s/U*) is also relatively large, although still much smaller than *L*. In practice, logarithmic parameters are never really large and corrections are important. Qualitatively similar behavior will arise as long as the population is in the multiple mutations regime with a large diversity of fitnesses in the population and the strong-selection weak-mutation regime so that growing subpopulations are only affected by fluctuations or mutations for a short time after they establish. To be quantitatively good, requires larger, but not unrealistically large, populations. Simulations in fig. 1 show that the predicted dependence of the speed on λ/*L* is quantitatively good with realistic parameter values: e.g. *N* = 10^12^, *s* = 10^−2^, 2*U* = 10^−4^ corresponding to *L* ≅ 23, *ℓ* ≅ 5.3 and thus *q* ≈ 9.

The logarithmic scaling of λ means that much of the benefit of sex can be obtained from very low mating rates. With these parameters, *r* ~ 10^−7^ would already yield a quarter of the maximum possible gain in speed (due to rapid reassortment but no recombination), while *r* ~ 2*U* = 10^−4^ would yield more than half. More importantly the oscillatory dynamics, as shown in fig. 4, are already very similar for these parameters to the deterministic asymptotic limit. And the period of oscillation in fig. 5 agrees quantitatively.

Note that although 10^12^ is a large population by standards of most laboratory evolution experiments, it is certainly not for all, e.g. the tabletop size MEGA plate has 10L of media and is capable of reaching a total of 10^12^ bacteria for typical cell densities of 10^8^/mL. [45]. And on scales of even a single human, it is less than the population sizes of the abundant gut bacterial species and the number of virions produced during the course of some viral infections [46].

### 9.4. Natural populations

Our model is not directly applicable to any real microbial populations, but it is most natural for the evolution of segmented RNA viruses with genomes divided into a number of segments that can reassort. To date, there are eleven families of segmented RNA viruses with the number of segments ranging from two to twelve [47]. For example, the bacteriophage *φ*6 has three segments and has been developed into an experimental system for both asexual and sexual evolution [48, 49]. Influenza viruses have six to eight segments. Nevertheless, we anticipate that many of the same features will apply. In a single host, or in a bacterial population, a viral population can be relatively well mixed without prominent spatial structure and co-infection rates, which are needed for reassortment, can vary widely. Thus the basic assumptions of the class of models we consider are reasonable.

The assumption that reassortment occurs but recombination within chromosomes does not—or at much lower rates—makes such models potentially applicable to chromids, or secondary chromosomes, found in an increasing number of species of bacteria. Chromids are hypothesized to derive from plasmids that have acquired essential genes from the primary chromosome [50]. They may retain the plasmid’s ability to transfer via conjugation. For example, *Pseudomonas syringae* pv. *lachrymans* has a recently acquired chromid that is self-transmissible via conjugation despite its large size of 1 Mb [51]. The possibilities of such large transfers of genetic material could alter the evolutionary dynamics of the species even if it occurs at very low rates and involves only better, rather than new, functions. For example, during the recent ecological differentiation of two populations of *Vibrio cyclitrophicus*, one of its two chromosomes swept within one population independently of the other chromosome, suggesting that reassortment played an important role [52].

### 9.5. Generalizations and extensions

How many of the features of our simple model obtain more generally? The scaling of the speed with λ/*L*? The sustained oscillations between mutation dominated and recombination dominated phases? Generalizations of the model can start to answer these questions and should be analyzable by a combination of methods used here and those developed by other. We outline a few of these and then discuss briefly the complications associated with richer— and more realistic—generalizations.

We have studied only a simplified model in which all mutations are considered to have the same selective advantage and there are no deleterious mutations. For large asexual populations, distributions of fitness effects have been studied and the primary results are that as long as the mutation rate spectrum, *μ*(*s*)ds to mutations with fitness effects *s*, falls off faster than exponentially, there is a predominant *s* (and a narrow range around this) that controls the evolution, with the rest being either too rare to matter, or effectively neutral [41, 40]. Inclusion of these effects into the two-chomosome model should be straightforward and result in a similar replacement of s and U by effective values that are determined primarily by *μ*(*s*).

Although usually phrased as an approximation of additive fitness effects of the mutations, the asexual model is far more general: it applies for a general fitness “landscape” (with arbitrary epistasis) as long as the statistical distribution of available mutations does not depend on the current genome. With recombination, however, combinations of interacting mutations are broken up and brought together in complex ways thus additivity is needed for the approximations used in almost all analyses of rapid evolution with multiple mutations to be valid. (A counter example, albeit without new mutations, is Neher and Shraiman [53].) In our model, however, all that is needed is additivity of the fitnesses of the two chromosomes and the requirement that the distribution of mutation effect sizes for both chromosomes remains unchanged during the evolution.

A natural generalization of the two-chromosome model is to multiple chromosomes that can reassort in some way, for example by exchange of a single one, or by grouping random combinations from two (or more) “parents”. With K chromosomes, the evolution speed will increase from *υ_a_*(*KU*) to *Kυ_a_*(*U*) (with *U* the beneficial mutation rate per chromosome) as r is increased from of order 1/*N* to of order s. Again, we expect the scaling parameter λ/*L* to primarily control the crossover. An important question—both here and more generally—is whether this *K*-chromosome model spontaneously oscillates. If it does, there should also be some *ℓ* dependence even in the asymptotic large-logs limit. To begin to address this, we can consider the simpler limit of *ℓ* → ∞ and λ/*L* → 0 — i.e. just barely enough reassortment to matter — discussed in section 7.1. For two chromosomes, the fittest offspring are from parental lineages that, since their last reassortment, mutated only on the X or only on the Y chromosome. For reassortment of *K* chromosomes between two parents, the fittest offspring are similarly from parental lineages that accumulated mutations on complementary sets of chromosomes. This implies that the maximum offspring fitness would increase at twice the speed of the mean for any *K*. Surprisingly, the dynamics of the resulting infinite *ℓ* oscillations are independent of *K* for λ/*L* = 0 and have speed *υ* ≈ 1.3 *υ_a_*. Understanding how the *K* > 2 oscillation dynamics change with *ℓ* and λ is left to future work. As an illustration of the behavior, simulation results for *K* = 3 are shown in fig. D.15: these show that the dynamics are similar to the *K* = 2 case for both low and intermediate mating.

An interesting question for the *K* chromosome model is how the dynamics change when approaching the sexual limit for which the speed approaches *Kυ_a_*(*U*). Already in the two chromosome case, we found a qualitative change in the dynamics when chromosomes can undergo multiple reassortments before accumulating more mutations. A regime in which such multi-reassortment processes dominate the dynamics was considered by Neher et al. [29]. They studied the sexual dynamics of models essentially equivalent to *K* → ∞ reassortment models but with the total mutation rate *μ* = *KU* fixed as the large *K* limit is taken so that new mutations always occur on different “chromosomes”. They were only able to analyze the dynamics for 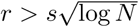 finding that the speed goes as *υ* ~ *r*^2^log(*Nμ*) up to other logarithmic factors that depend on the particular reassortment processes. In contrast to the two-chromosome model for *r* ≪ *s*, mutations that arise in the bulk of the fitness distribution contribute substantially: multiple reassortments enable them to combine onto better genomic backgrounds and eventually to the nose. This enables many mutations to segregate simultaneously (although almost all are still wasted, as *υ* ~ log(*Nμ*) ≪ *Nμ*). The dynamics of each new mutation proceeds in the distribution of fitness backgrounds of the other segregating mutations and the new mutant lineages only feedback to affect the earlier mutations when their frequency has risen enough in the population that the distribution of the new mutation over the fitness backgrounds has become essentially deterministic. But as *r* decreases, the fluctuations of the nose of the fitness distribution become large enough that this approximate independence of each mutation breaks down. Accumulation of multiple mutations on the same genome before reassortment then becomes important. Once this occurs, the details of the distribution of relatedness becomes essential. It is not known how the large *K* models behave for *r* ~ *s* or *r* ≪ *s*, but we expect that the crossover of the speed to asexual should again be only logarithmically dependent on *r*. Understanding the underlying complex dynamics, which will involve subpopulations that accumulate mutations and mate in anomalously rare ways, is a real challenge for future research.

The dynamics of facultative sexual populations with very low rates of mating but a large number of crossovers when they do mate—completely destroying linkage—was studied by Rouzine and Coffin [33, 34, 35]. Although they consider only purifying selection on preexisting deleterious variation with no new mutations, there is still a range of times in which the dynamics is well approximated by a steadily moving fitness wave. Their analysis exhibits some general features similar to ours: a dependence on logarithmic parameters equivalent to our log(*Nr*)/log(*Ns*) in the crossover regime, and a cycle of recombination and growth that advances the nose, simpler but loosely analogous to the wing-supporting cycle in our steady state solution. But they treat the complex correlations induced by common ancestry in a relatively simple manner [35] which is only a crude approximation.

The most interesting direction is moving away from the non-recombining chromosome models towards more realistic recombination processes. One example is facultatively sexual organisms that occasionally mate and when they do so, the chromosomes recombine with a few crossovers, in addition to reassorting. As each of the long segments that remain linked will have evolved asexually for some time before recombining, this has some features that are crudely similar to the *K*-chromosome purely-reassorting model. But the crossovers occur at different positions in different matings. Thus the many possible segments of the chromosomes in the many possible individuals in the population that could be recombined together need to be kept track of. Whether this can be done in some approximate way in terms of many effective non-recombining chromosomes— loosely analogous to the treatment of asexual segments of chromosome at moderate recombination rates by Weissman and Hallatschek [31], Neher et al. [30], Weissman and Barton [32]—is unclear. In any case, this certainly represents an important and challenging direction for future research.

Another challenging direction is applicable to bacteria, most of which primarily exchange small segments of chromosomal DNA. Of course, new functions can be acquired as single genes or whole operons. But even homologous recombination of “uninteresting” segments can contribute much more to the fitness than those they replace because of accumulation of beneficial mutations: this is the natural generalization of the evolutionary processes we have analyzed. Again, it is plausible that the dynamics could be analyzed in terms of segments that are effectively like short reassorting chromosomes—loosely analogous to the *K*-chromosome model for some effective *K* but with exchange of only one chromosome at a time.

For all of these extensions, the most interesting observable features may well be the statistics of diversity induced by the dynamics. As for rapidly evolving asexual populations [27, 28], these will reflect the crucial but invisible dynamics of the very low frequency subpopulations that drive the dynamics, i.e. the “nose that wags the dog” [44].

## Appendix A Two Chromosome Asexual

Under the assumptions discussed in section 5, there will be a steady state front that moves with a constant speed *υ*. In the continuous fitness approximation, the absolute fitness of the front will take the form *F*(*w,t*) = *υt* + *f*(*w*) with *w* ≡ *X* − *Y*. A new subpopulation at the front will be fed by two subpopulations that were previously at the front. These two subpopulations will have the same absolute fitness *F* = *υτ*_1_ + *f*(*w*_1_) = *υτ*_2_ + *f*(*w*_2_), where the τ’s are their respective establishment times. The difference in *w* values is *w*_2_ − *w*_1_ = 2*s* because *w*_2_ = (*X*_1_ + *s*) − (*Y*_1_ − *s*). Therefore

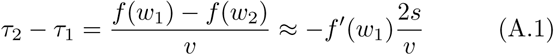

with 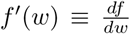. The newly established subpopulation has fitness *F*+*s* = *υτ*_est_+*f*(*w_i_*), where *w_i_* = *w*_1_+*s* = *w*_2_ —*s* is the intermediate w value. Thus its establishment time must be

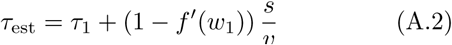

Using eqs. (A.1, A.2) and the establishment time from the feeding process found in eq. (15) gives a differential equation for the steady state asexual front

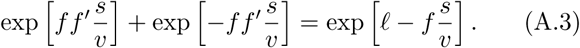

**Figure A.9:**
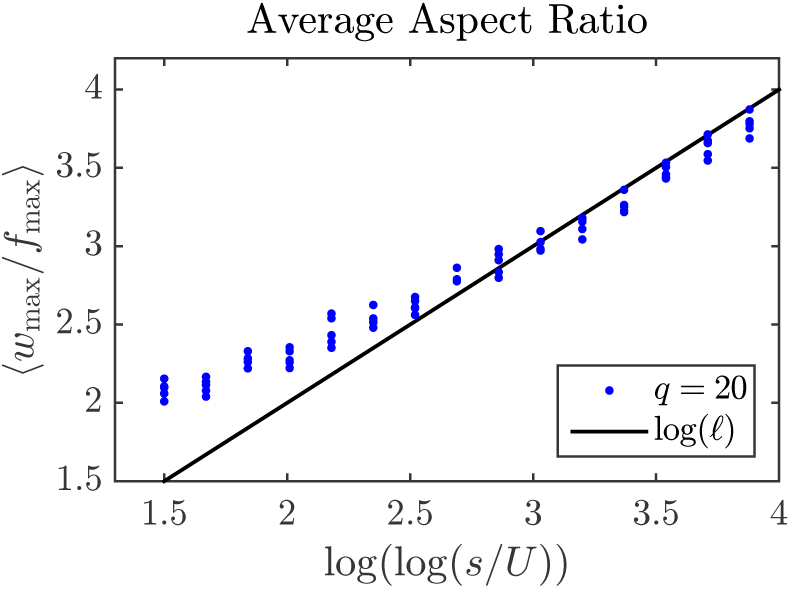
Comparison of the stochastic simulation results for the aspect ratio averaged over time for two different values of *L* ≡ log(*Ns*) and a range of *ℓ* ≡ log(*s/U*) values. For large *ℓ* the simulations are in good agreement with the asexual steady state calculation.

Rescaling fitness variables *f* and *w* according to eq. (9) shows that *ℓ* controls the overall shape:

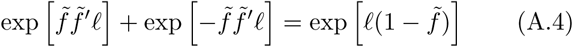

Numerical solutions to this equation are shown fig. 2.

There are two regions of the front. The region near the nose (which we choose to be at *w* = 0) experiences roughly equal mutational feeding by the two parent subpopulations, i.e. τ_1_ and τ_2_ are nearly equal. The outer regions called the wings have feeding dominated by a single subpopulation, so either τ_1_ or τ_2_ is much earlier. The transition between the two regions sets the overall width of the fitness distribution which varies with the mutation parameter *ℓ* = log(*s/U*).

The nose region with two-sided feeding has roughly the same width for different values of *ℓ*, but the fitness drop from the nose goes as *O*(1/ *ℓ*). To see this, expand eq. (A.4) to lowest order in 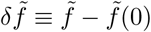:

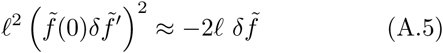

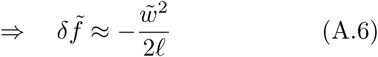

This solution breaks down when 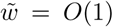 and 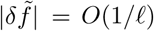.

The width of the wing depends on its starting fitness. The X wing 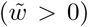 starts approximately at the end of the nose region, with 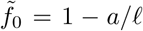 and 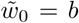 for some order one constants *a, b* with which we can roughly match the nose and wing regions. When 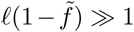 either 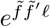 or 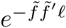 in eq. (A.4) must be much larger than the other. Therefore

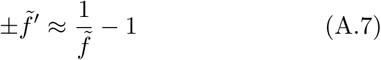

where the sign depends on which mutation wing we are considering. The solution for the X (*w* > 0) wing starting with 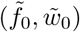 is

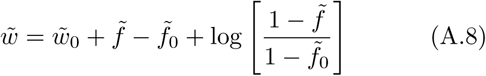

The rightmost edge of the distribution is at 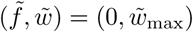 so plugging in for 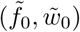 we find

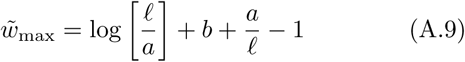

so 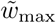, the half-width of the distribution, approaches log(*ℓ*) + *c* for large *ℓ*. In terms of the original parameters this is 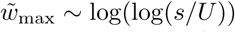) which grows very slowly with the mutation timescale 1/*U*. Fig. A.9 confirms this scaling in the stochastic simulations by considering the “aspect ratio”, *W*_max_/*f*_max_, where *f*_max_ = *qs*[1 − 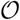(1/ *ℓ*)]. This ratio is the same as that for the rescaled quantities: i.e. equal to 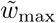.

It would appear that this analysis depends crucially on the continuum approximation for fitnesses which for fixed large *q* will break down for rescaled fitness differences of order 1/*q*. Yet the fitness thickness of the nose regime was inferred to be only of order 1/ *ℓ* suggesting breakdown when *ℓ* > *q*. But the results shown in Fig. A.9 appear to indicate that the results are valid even far into this regime. Although we have not analyzed this in detail, it appears that the crucial property is that the establishment times for a given fitness vary slowly with *w*, even if the fitness steps from *Z* to *Z* + s are themselves very jerky as they are when *ℓ* ≪ *q*.

## Appendix B. Intermediate Mating

### *Appendix B.1.* Steady State

For mating rates λ/*L* ≳ 0.81, we are forced into an *Ansatz* for the steady state with the front divided into three regions: the nose region, the mutation wings, and “re-mating” regions between the nose and wings, as illustrated in fig. B.10. As in the low mating steady state, the nose region is the product of reassortment from both wings and therefore has the same shape 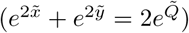. Due to the higher mating rates and corresponding faster speed, the nose region is supported by mutation wings that start further from the nose (larger 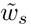) with lower initial fitness 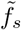 than the low mating case. The lower fitness wings can support a nose region of limited width. The gaps between the nose region and the wings must then contain subpopulations that establish due to reassortment (since they are not in the wings) and later contribute to reassortment after clonal growth, hence the name re-mating. The X re-mating region is the product of reassortment between the X wing and the Y re-mating region. In fig. B.10 we can follow the course of a particular X chromosome that begins at the X mutation wing start: it accumulates mutations and then grows in copy number. Some copies reassort and establish in the Xside re-mating region. They again grow in copy number and some reassort and establish in the other Y re-mating region. These chromosomes have thus undergone two reassortment events. They will not reassort again because their *X* fitness is now too low to establish at the front.

**Figure B.10:**
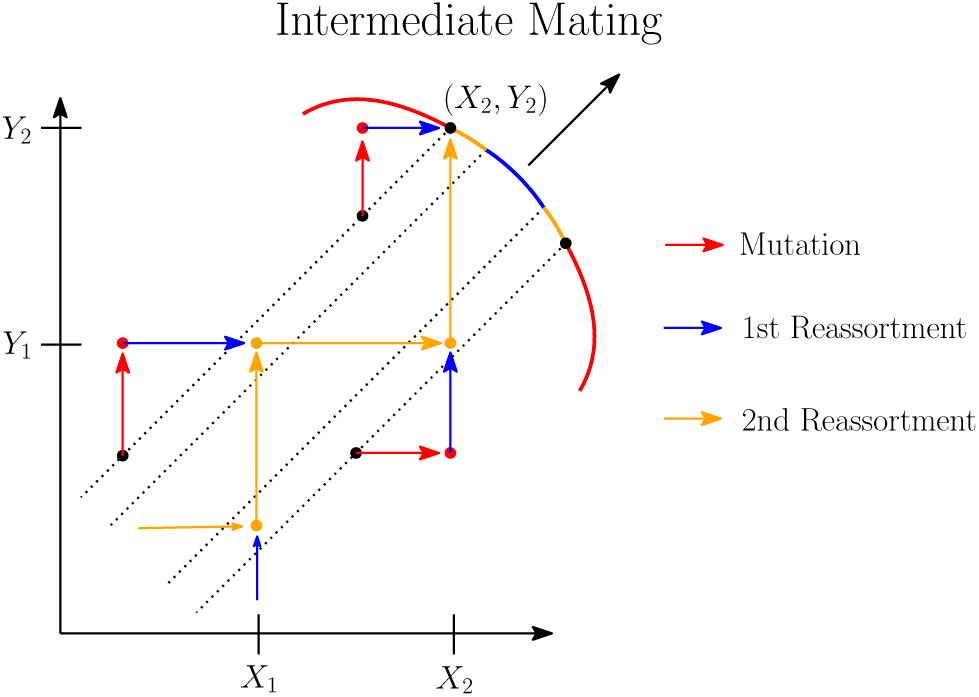
Schematic of the steady state dynamics for intermediate mating rates. There are three regions of the front: the mutation wings (red), the nose region (blue), and the re-mating regions (orange). The subpopulations in the nose region establish due to reassortment from the two mutation wings. Subpopulations in, e.g, the X-side re-mating region are established by reassortment of a chromosome from the Y re-mating region and a chromosome from the X mutation wing. The remating subpopulations both establish by reassortment and contribute to reassortment. The diagram shows the series of reassortments that ultimately support the wing start. Consider an X chromosome at the wing start. It will accumulate mutations (horizontal red arrow) and reach a fitness of *X*_2_ (bottom right). It grows clonally for a time and then successfully reassorts (vertical blue arrow) to the X re-mating region. It grows again and then reassorts (vertical orange arrow) to the Y wing start at (*X*_2_, *Y*_2_).

As in the low mating regime, the long term dynamics are controlled by an essential wing-supporting cycle that allow a chromosome to persist indefinitely by accumulating mutations and reassorting back to the wing start again and again (see fig. 3). For the Y-wing, this cycle supplies the Y chromosome. A new complication is that the *X* chromosome for the Y wing start does not simply come from the other mutation wing. Fig. B.10 traces back some of the reassortment events needed to eventually support the wing start. The X chromosome for the Y-wing start comes from a point in the opposite (X) re-mating region. This point is itself also supported by a point in its opposite (Y) re-mating region, and so on. There is an infinite sequence of points supporting each other. Backwards in time, the sequence moves inward from the wing start at the edge of the re-mating region and converges to a fixed point in its interior. So the dynamics further and further back in time depend on a smaller and smaller region around the fixed point. This steady state could only be approached from initial conditions by dynamical transients continually building up the region around the fixed point.

It is possible to determine the speed of the intermediate mating *Ansatz* without explicitly solving for the whole front *f*(*w*). Given the fitness and slope, (*f*(*w*),*df*/*dw*), for a point in the re-mating region we can solve for the points that support it by reassortment. So from the wing start at 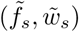, we can find the sequence of points backwards in time. For the correct set of parameters (the speed v and the wing start at fixed mating rate λ/*L*) the sequence will converge smoothly to a fixed point. For an incorrect set of parameters, the sequence will diverge or not matchup smoothly. By adjusting the parameters we can thereby determine how v depends on λ/*L*.

First we describe how reassortment, in general, constrains the shape of the front 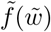 by relating distant points. Reassortment is controlled by the total number of individuals with a given fitness of each chromosome, *n*(*x*) = ∑_*y*_ *n*(*x, y*) (and visa versa). Because of exponential growth, different subpopulations *n*(*x, y*) differ by orders of magnitude, so (to the needed logarithmic accuracy) the sum is dominated by its largest term: *n*(*x*) ~ max_y_ *n*(*x, y*). As a subpopulation initially at the front grows, its fitness decreases from 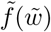 to 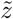 in a time 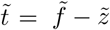 so its size is

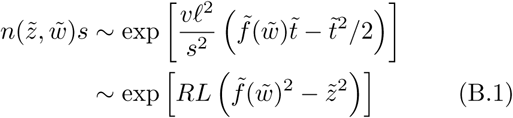

with 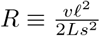 Let *m* be the fitness of the maximum over *y* for a given 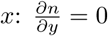 gives 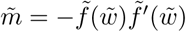 so that

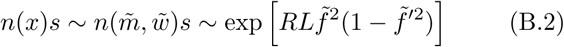

The subpopulation at 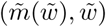 is the dominant supplier for X chromosomes with relative fitness 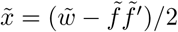.

Reassortment connects the subpopulation 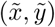 estab-lishing at the front and the two parent subpopulations, 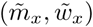 and 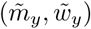, which supply the X and Y chromosomes to 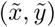. Of course the parent subpopulations had previously established at the front-at 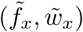 and 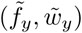—before growing. The reassortment constraint, explained in section 5.1, is that the influx of reassorters 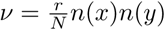 satisfies log *ν* = 0, which becomes

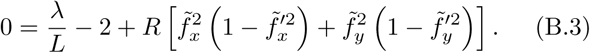

after using eq. (B.2). We also need 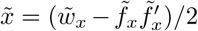 and 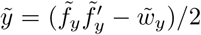 to properly connect the corresponding points.

Next, we apply the general reassortment equation, eq. (B.3) to the intermediate mating *Ansatz*. The boundary conditions for the re-mating region can found in terms of the wing start parameters. One edge of the X re-mating region is at the X wing start, 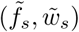, and has a slope of 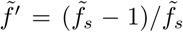, from eq. (A.7). The other edge is the endpoint of the nose region, which is the furthest point that the Y wing can support via reassortment. Therefore the nose region endpoint at 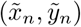 is supplied Y chromosomes from a subpopulation growing directly from the Y wing start at 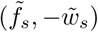. So 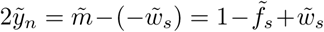, and 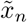 and the slope can be inferred from the nose region shape, 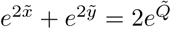

**Figure B.11:**
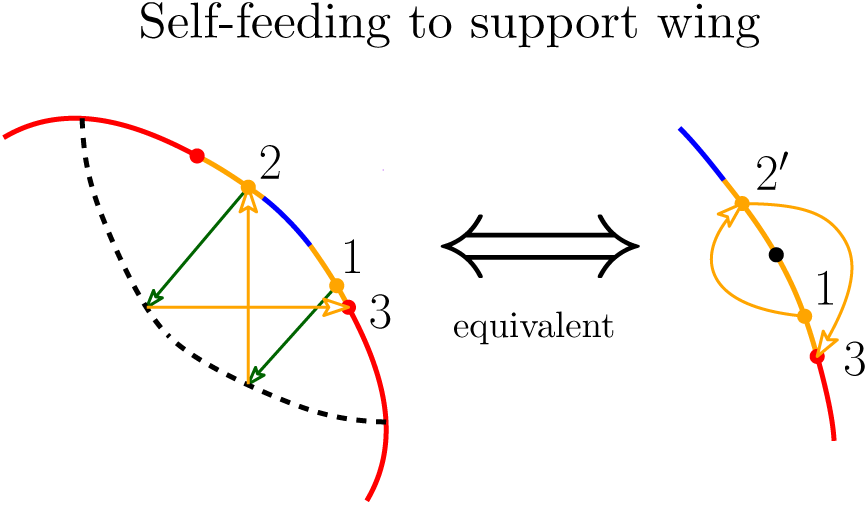
A lineage starting in the X re-mating region (at point 1) grows for a time (green arrow) and reassorts (orange arrow) to the Y re-mating region (at point 2) and then grows and reassorts again to point 3, shown to be the wing start. Since the re-mating regions have the same shape up to a reflection across *x* = *y*, we can reinterpret this reassortment process as a map from one re-mating region to itself, as shown in the right diagram with the corresponding points labeled. Point 2’ is the reflection of point 2. The map expands outward so there is an unstable fixed point (black dot) somewhere in the interior of the re-mating region. The dashed black curve on the left denotes the line of subpopulations with maximum size among subpopulations with the same X or Y chromosome fitness.

The contribution to eq. (B.3) from the mutation wing can be found simply. From eq. (27), the X wing has fitness 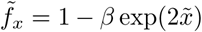 with 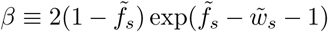 as a useful combination of parameters. Since 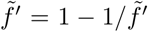, the wing contribution is

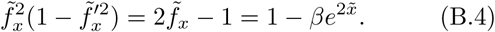

By plugging the wing contribution into eq. (B.3), we effectively account for one of the three points and obtain a general relation between two points, both in the re-mating region.

The parent subpopulation in the Y re-mating region with 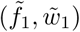 supports the establishment of the point 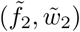 in the X re-mating region. The child subpopulation receives an X chromosome with fitness 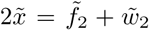 from the mutation wing. Equation (B.3) becomes

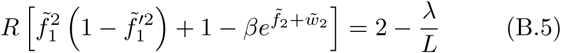

with the additional constraint that the Y chromosomes from 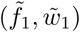 have the correct fitness:

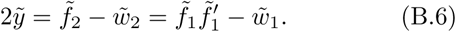

The slope of the front at the child subpopulation can be found by differentiating eqs. (B.5) and (B.6) to obtain?

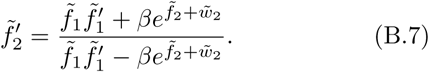

Together these three equations determine the front at 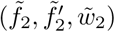, in the X re-mating region as a function of 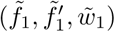 in the Y region.

Since we assume that the *Ansatz* is symmetric, we can modify eqs. (B.5) to (B.7) to involve two points in the X re-mating region using the substitution: 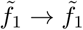, 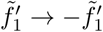, and 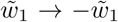. The modified equations are a map from one point in the re-mating region to the another. It can be shown that this reassortment map has a fixed point in the interior of the region and application of the map expands points outward from the fixed point. Figure B.11 illustrates how reassortment in the re-mating region can be reinterpreted as an expanding map.

To solve for the steady state, we must use the inverse of the reassortment map to move inwards from the edges to the fixed point. For the correct set of 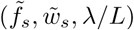 values, the inverse map trajectory reaches the fixed point smoothly, but for incorrect values the trajectories diverge or intersect at a kink. So the correct values can be found via the shooting method used to solve boundary value problems: simply vary the parameter values until a solution is found. As for low mating rates, the nose fitness can be solved for in terms of λ/*L* using eq. (32), which determines the speed via 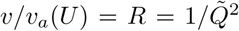. For the steady state solution, the re-mating region expands and the nose region shrinks as the mating rate increases. For the *Ansatz* to be valid we must check that the nose region exists. This amounts to checking whether 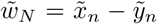 is greater than zero. This condition fails for λ/*L* ≥ 0.99 suggesting that new *Ansatzes* are needed for mating rates approaching *r* ≲ *s* that involve additional reassortment events.

### *Appendix B.2.* Oscillations

Comparison of the intermediate oscillations in fig. B.12 and the low mating oscillations in fig. 4 show notable differences in the establishment history of the current population: At low mating rates subpopulations established during a single reassortment phase are present, but at intermediate rates the current population contains subpopulations from two different reassortment phases coexisting. This difference is due to a change in the dynamics of the oscillations. Fig. 5 shows how the period in the deterministic simulations depends on the mating rate. As the mating rate increases, there is a discontinuous period-halving bifurcation at λ/*L* ≈ 0.8, which essentially coincides with the transition point between the low and intermediate mating-rate steady states. When the period halves, two reassortment segments can “fit” within the current population instead of just one. (The simplest scenario is that decreasing λ from the intermediate mating rate regime leads to a sub-critical—and hence discontinuous— period doubling bifurcation, while increasing it from the low mating rate regime leads to the disappearance of the cycle at a saddle-node bifurcation. This makes the transition between the two regimes hysteretic as is, indeed, observed in the deterministic simulations.)

As for the low mating oscillations, small regions of the front at the end of the reassortment phase produce the mutation wings important for reassortment and are analogous to the wing starts for the intermediate steady state. The set of reassortment events that support these small regions resemble the intermediate wing-supporting cycle in fig. B.10. Again the intermediate mating oscillation appears as if one cycle from the steady state has strengthened to dominate the others. The correspondence between cycles from the steady state and the oscillations is qualitative but not necessarily quantitative since the oscillation cycles must satisfy all the delayed feedbacks. The difference between low and intermediate mating oscillations is an additional feedback due to the secondary reassortments. Consider the marginal case with λ/*L* at the period halving transition point. Initial transients appear as low mating oscillations with a long reassortment phase. The crucial difference is that subpopulations established at the beginning of the reassortment phase also contribute to re-assortment by the end of the phase. This is the start of an extra reassortment phase and creates an additional feedback: the shape at the end of the (long) reassortment phase depends on the beginning of the reassortment phase. (For low mating oscillations the shape at the end depends only on the previous reassortment phase). The additional feedback eventually breaks the long reassortment phase into two separate phases. The jumping of the mean is now “interlaced” between cycles: one reassortment phase ends due to the mean jumping caused by the previous reassortment phase, as can be seen fig. B.12. The period halving point roughly coincides with the low-to-intermediate mating transition because both happen when the mating rate is large enough that a subpopulation established by reassortment can later contribute to reassortment.

At the upper end of the intermediate mating regime where *r* → *s* (i.e. λ/*L* → 1) the size of the mean jumps decays to zero and the reassortment regions become larger and begin to overlap. This behavior smoothly approaches the obligate sexual steady state, which always has establishment by reassortment and does not exhibit oscillations in the deterministic limit. For intermediate mating rates, the nose always has some influx of individuals due to reassortment—as seen in fig. B.12—although there are still clear reassortment and mutation phases. In the obligate sexual limit the two chromosomes become completely unlinked, so the distribution is a product of the asexual fitness distributions of each chromosome. The oscillation period, shown in fig. 5, becomes close to the nose-to-mean time, τ_nm_, of the sexual limit. Since the speed is twice as fast as the asexual limit, the time for the mean to advance one nose-length is half as long, i.e. τ_nm_ ≈ 0.5 *ℓ*/*s*. Thus for λ → *L*, the oscillation period approaches the time for variations in the speed of the nose to cause change in the speed of the mean. Note that this is not the nose-to-mean time for the distribution of one of the almost-independent chromosomes, which would be twice as long. Not surprisingly, the coupling between the chromosome distributions is still important for the oscillations.

**Figure B.12:**
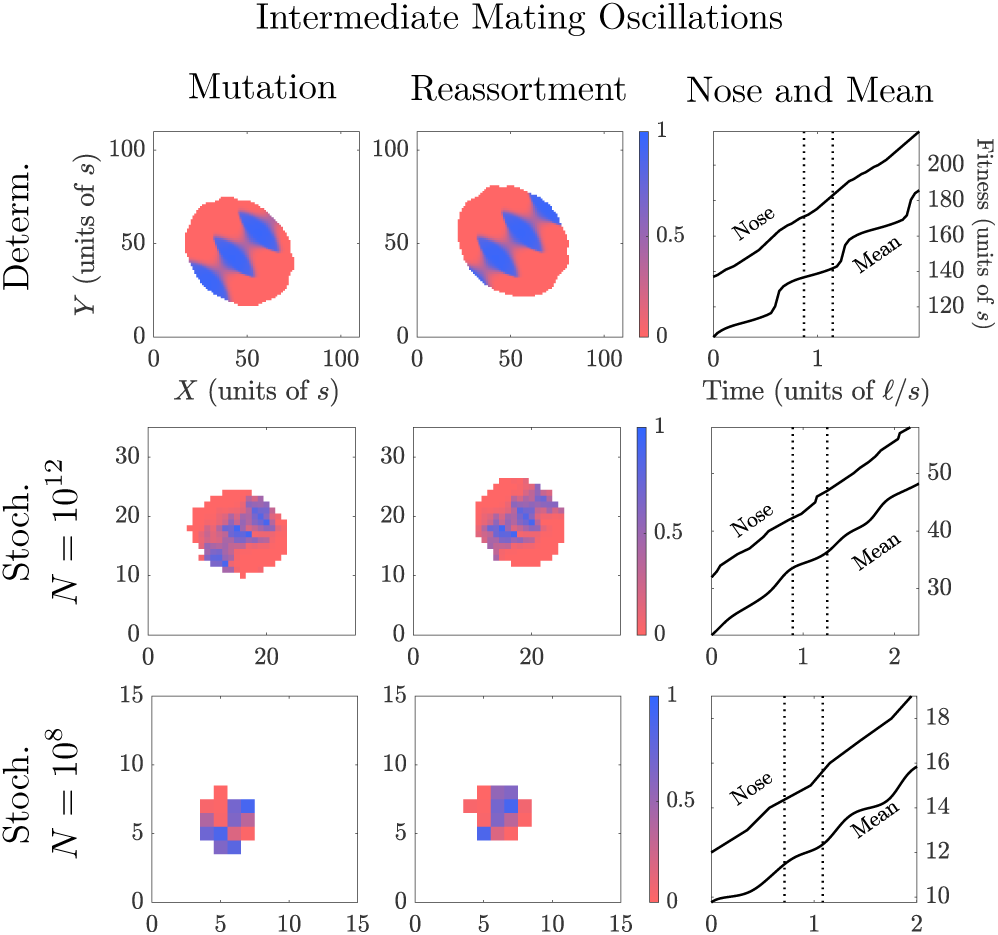
Oscillation cycles illustrating the fitness distribution during the mutation phase (left column) and the reassortment phase (middle column) which occur during each cycle; the top row is for deterministic dynamics. All the non-zero subpopulations are shown with their color indicating the fraction of new individuals due to reassortment (bluer) or mutation (redder) when establishing. The oscillation dynamics are shown for intermediate mating rates with λ/*L* ≈ 0.8. In this regime, the current population includes subpopulations established in two reassortment phases, instead of only one phase for the low mating oscillation shown in fig. B.12. (This is clearly seen even though the intermediate rate used is on the border of the two regimes.) The righthand column shows the speed of the nose and mean, dashed lines corresponding to the times of the snapshots shown. The nose speed increases during the reassortment phase and, through exponential growth of the prior nose populations, the effects of this are sharpened into a jump in the mean fitness roughly a time *ℓ*/*s* later. At intermediate mating rates, another reassortment phase occurs within the timespan required for this exponential growth. The stochastic simulations have *N* = 10^12^, *s* = 10^−2^, 2*U* = 10^−-4^, *r* = 10^−-4^ (*q* ≈ 9, *ℓ* = 5.3) and *N* = 10^8^, *s* = 0.03, 2*U* = 10^−6^, *r* ≈ 2 × 10^−3^ (*q* ≈ 2.7, *ℓ* ≈ 11). These agree qualitatively and semi-quantitively with the deterministic simulations which are valid in the continuous (large *q* ≡ 2*L*/*ℓ*) limit.

## Appendix C. Crossovers and deterministic-stochastic comparisons

### *Appendix C.1.* Crossover to asexual limit

The analysis of the reassortment steady state in eq. (34) shows that the growth of populations from the center of the front yields a fitness distribution near the mean that is approximately gaussian with an aspect ratio, 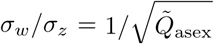, that approaches unity in the asexual limit for large *ℓ*. But the asexual steady state is far from symmetric. Using the results from Appendix A, a similar calculation finds an aspect ratio 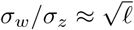. This implies that reassortment is a singular perturbation in the asymptotic limit, meaning that the shape of the front changes rapidly for a small increase in λ/*L*. The curvature of the front at the nose changes quickly over a narrow range of mating rates such that reassortment is unable to establish populations at the front for small λ/*L*.

The crossover from asexual is complicated by the *ℓ* dependence of the asexual steady state which has a speed of *υ_a_*(2*U*) = 2*L*/(*ℓ* − log2)^2^. The steady state dynamics are asexual until reassortment first results in establishment at the nose. We can derive how this critical reassortment rate depends on *ℓ*. A subpopulation with fitnesses 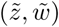 has a size

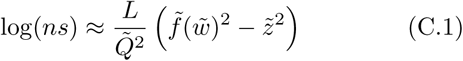

with 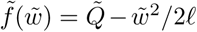 for the asexual front, as derived in eq. (A.6). The subpopulations feeding the nose are located at 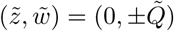 up to 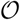(1/ *ℓ*) corrections. From the requirement for establishment (*rn_x_n_y_*/*N* ~ 1) we find that reassortment first matters in the deterministic approximation when λ/*L* = 2/ *ℓ*.

The reassortment steady state solution derived in section 6.1 represents the *ℓ* → ∞ limit. This solution has the correspondingly correct asexual speed *υ_a_*(*U*) = 2*L*/ *ℓ*^2^ and first deviates from asexual at λ = 0. Corrections for finite *ℓ* must therefore produce a solution with the correct asexual speed, *υ_a_*(2*U*), up to λ/*L* = 2/ *ℓ*. For finite (but large) *ℓ*, the mutation wing is described by the full differential equation for the asexual front, eq. (A.4). As explained in Appendix A, finite *ℓ* corrections are only relevant for fitnesses 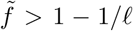. Since the wing start fitness 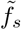 must be less than the nose fitness 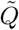, we can estimate that the corrections become significant only when 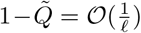 which as for the two-chromosome asexual corrections also occurs for 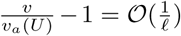. For speeds below this threshold, corrections help the solution asymptote to the correct asexual limit for finite *ℓ*. For speeds above this threshold, finite i effects are negligible and the solution matches the infinite *ℓ* steady state derived in section 6.1.

### *Appendix C.2.* Crossover of oscillation dynamics for finite ℓ

The speed curves in fig. 1 show considerable dependence on *ℓ* in the low mating regime. The changes of the speed are predominantly due to changes in the jump size of the mean fitness, as seen in the plots of the period and jump size in fig. 5. The size of the mean jump is determined by the nose speed at the start of the reassortment phase, which depends on when reassortment interrupts the mutation phase. In the absence of mating the population would eventually approach the asexual steady state. The oscillation dynamics for general *ℓ* and λ interpolate between two extremes, when either the population is close or far to the asexual steady state when reassortment starts.

The infinite *ℓ* limit described in section 7.1 is always far from the steady state (indeed no steady state formally exists in this limit). The front is an expanding flat line that advances at speed roughly *υ_a_*. The line of subpopulations with size, *n_c_*, sufficient to reassort to the nose trails behind the front, also advancing at roughly speed *υ_a_*, like the ridge in fig. 7. The *X*-most point of the *n_c_* line moves purely in the *X* direction at speed *υ_x_* = *υ_a_*, and similarly for the *Y* -most point. During the reassortment phase, this implies that the nose speed is *υ_n_* = *υ_x_* + *υ_y_* = 2*υ_a_*. According to eq. (37), the nose-mean ratio *β* = *υ_n_*/*υ_a_* ≈ 2 results in a jump in mean fitness of size 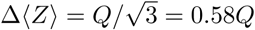. This regime applies for large *ℓ* (so that the approach to steady state takes a long time) with moderate λ/*L* > 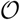(1/ *ℓ*) (so that the mutation phase is not too long). In fig. 5, the large *ℓ* jump sizes plateau to 0.58Q up to 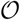(1/ *ℓ*) corrections.

In the opposite regime—for fixed *ℓ* as the speed is just rising from the asexual speed—the population is close to steady state when reassortment starts. This happens for λ/*L* ≈ 2/ *ℓ* when reassortment is just enough to matter for the asexual steady state, as derived in the previous appendix section. In the steady state, the whole population advances at *υ_a_* in the *X* = *Y* direction so the *X*-most point with *α_c_* moves at only *υ_x_* = *υ_a_*/2 in the *X* direction. The reassortment nose speed is therefore *υ_n_* =*υ_x_* + *υ_y_* = *υ_a_*, implying (from eq. (37)) a vanishing jump size: this is a marginal case. Small values of *ℓ* are more similar to the steady state because the 1/ *ℓ* effects induce curvature of the front earlier in the mutation phase. So for small *ℓ* the fitness distribution and the speed remain in the crossover regime between the steady state and the infinite *ℓ* limits for a greater range of λ/*L* values in fig. 5.

For intermediate mating rates, the speed curves in fig. 1 are *l* independent. This likely happens because the lineages important for reassortment descend from points that start the mutation process in the single parent regime. Then *ℓ* can be scaled away as in the reassortment steady state analysis.

## Appendix D. Additional figures

**Figure D.13:**
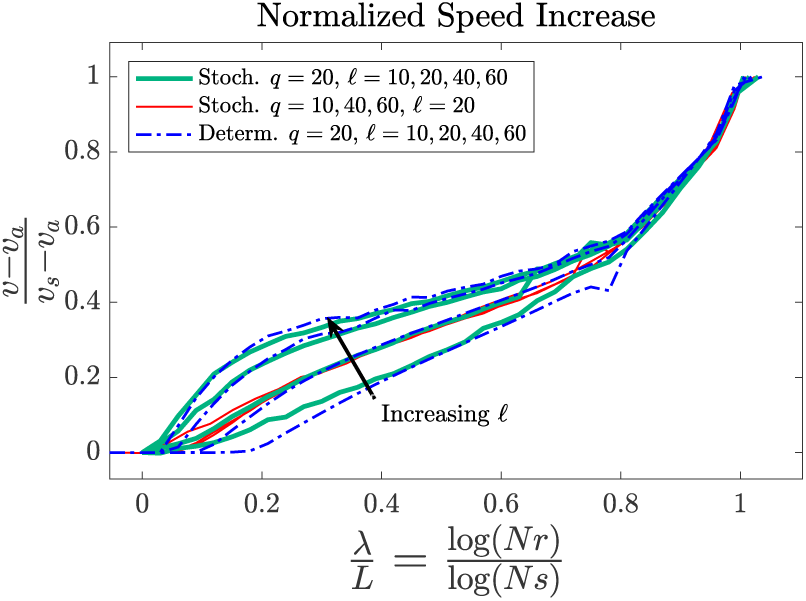
Speed comparison between deterministic (blue) and stochastic simulations (teal) for a range of *ℓ* ≡ log(*s/U*). Smaller *ℓ* values appear lower in the plot. As discussed in section 5.1, the first speedup due to reassortment is at λ/*L* = 2/ *ℓ* for the deterministic model. Reassortment affects the stochastic dynamics for smaller λ/*L* due to fluctuations in the width of the asexual distribution. The red curves, which are stochastic simulations with fixed *ℓ* but different *q*, overlap greatly and show very little dependence on *q* except when stochastic effects are important for λ/*L* < 2/ *ℓ*.

**Figure D.14:**
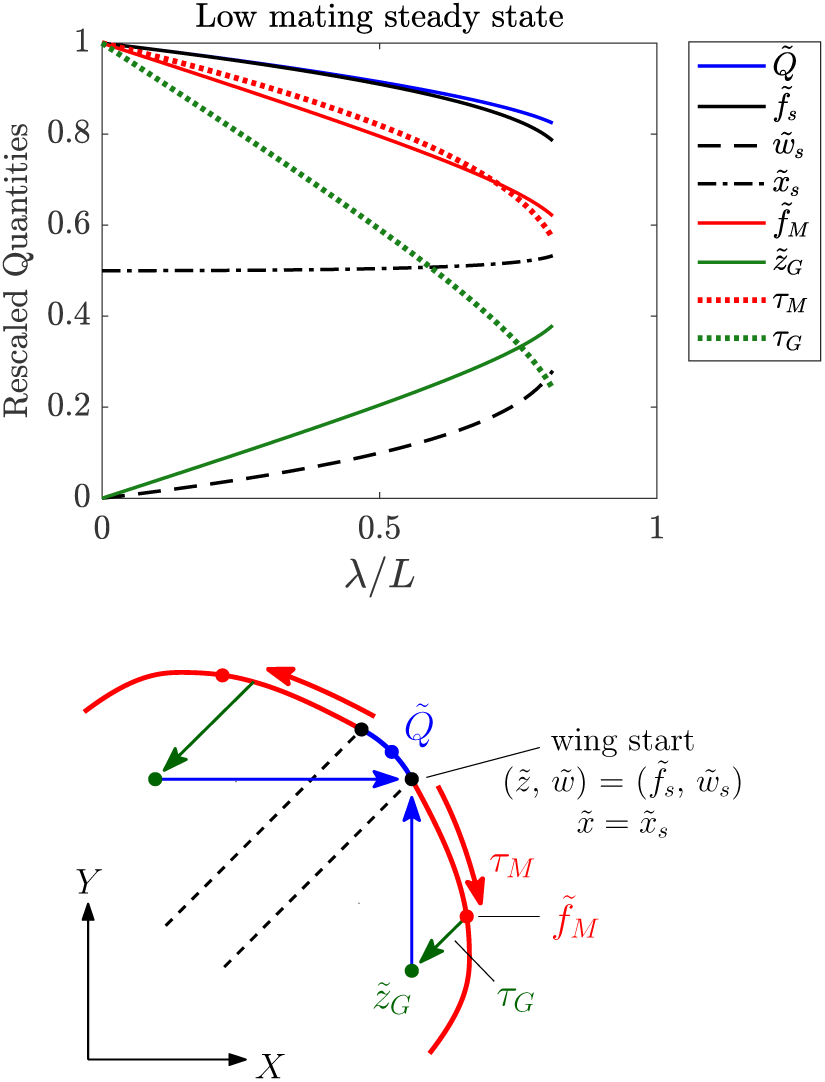
The dependence on reassortment rate of low-mating steady-state quantities, rescaled according to eq. (9). The diagram on the left illustrates the quantities plotted. These include the nose fitness 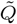, the fitnesses for the wing start: fitness 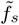, transverse fitness 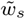, and X chromosome fitness 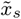. For the wing-supporting cycle, the fitnesses after mutation, 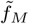, and after growth, 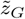 together with the times for mutation, τ_*M*_, and growth, τ_*G*_, are also plotted.

**Figure D.15:**
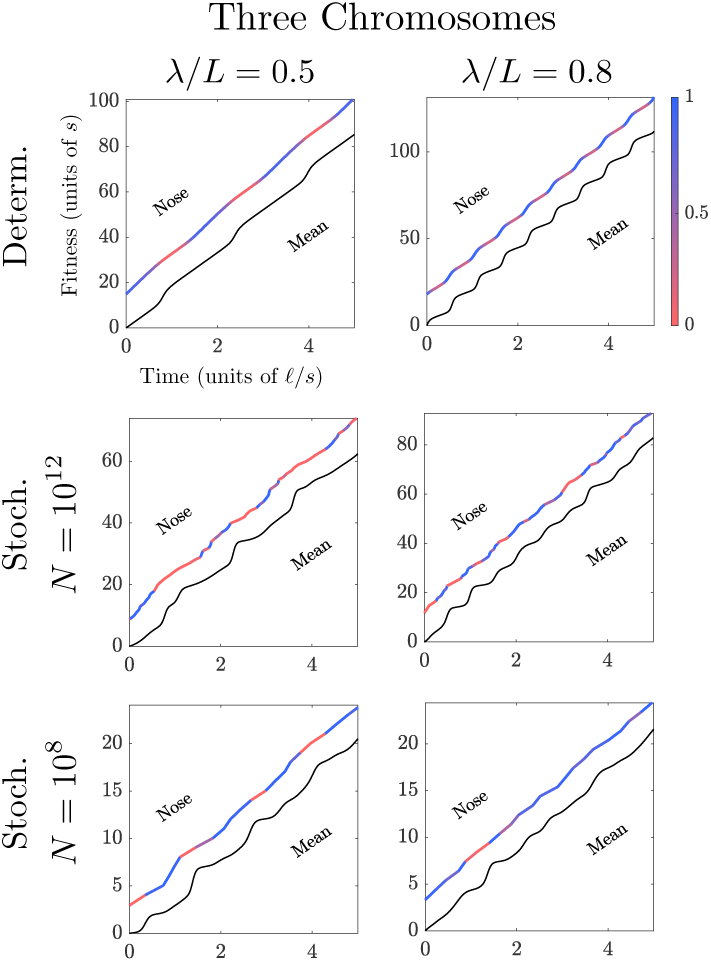
The nose and mean fitness trajectories for the three chromosome (*K* = 3) model. The nose trajectory is colored by whether establishment is due to a greater influx of reassorters (bluer) or mutants (redder). Sexual reproduction is implemented using the “communal” model of Neher et al. [29] in which each chromosome is sampled separately from the population, so the reassorted population is 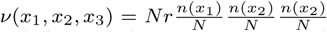. The oscillation dynamics at low (λ/*L* = 0.5) and intermediate (λ/*L* = 0.8) mating rates are similar to the two chromosome case shown in figs. 4 and B.12. The realistic parameter values used are the similar to the two chromosome plots: *N* = 10^8^, *s* = 0.03, 3*U* = 10^−6^ and *N* = 10^12^, *s* = 0.01, 3*U* = 10^−4^.

## Appendix E. Acknowledgements

We would like to thank Atish Agarwala for helpful and enjoyable discussions. MTP is supported by William R. and Sara Hart Kimball as a Stanford Graduate Fellow and by the National Science Foundation via a Graduate Research Fellowship, DGE-114747. This research was supported by the National Science Foundation via PHY-1305433 and PHY-1607606.

